# Elucidating the activation mechanisms for bifurcation of regulatory and effector T cell fates by multidimensional single cell analysis

**DOI:** 10.1101/280818

**Authors:** Alla Bradley, Tetsuo Hashimoto, Masahiro Ono

**Author notes:** Address correspondence and reprint requests to Dr Masahiro Ono, Department of Life Sciences, Sir Alexander Fleming Building, Imperial College Road, South Kensington, London, SW7 2AZ, United Kingdom.

## Abstract

In T cells, T cell receptor (TCR) signalling initiates downstream transcriptional mechanisms for T cell activation and differentiation. Foxp3-expressing regulatory T cells (Treg) require TCR signals for their suppressive function and maintenance in the periphery. It is, however, unclear how TCR signalling controls the transcriptional programme of Treg. Since most of studies identified the transcriptional features of Treg in comparison to naïve T cells, the relationship between Treg and non-naïve T cells including memory-phenotype T cells (Tmem) and effector T cells (Teff) is not well understood. Here we dissect the transcriptomes of various T cell subsets from independent datasets using the multidimensional analysis method Canonical Correspondence Analysis (CCA). We show that resting Treg share gene modules for activation with Tmem and Teff. Importantly, Tmem activate the distinct transcriptional modules for T cell activation, which are uniquely repressed in Treg. The activation signature of Treg is dependent on TCR signals, and is more actively operating in activated Treg. Furthermore, by analysing single cell RNA-seq data from tumour-infiltrating T cells, we revealed that FOXP3 expression occurs predominantly in activated T cells. Moreover, we identified FOXP3-driven and T follicular helper (Tfh)-like differentiation pathways in tumour microenvironments, and their bifurcation point, which is enriched with recently activated T cells. Collectively, our study reveals the activation mechanisms downstream of TCR signals for the bifurcation of Treg and Teff differentiation and their maturation processes.

## Introduction

T cell receptor (TCR) signalling activates NFAT, AP-1, and NF-κB (1), which induces the transcription of Interleukin (IL)-2 and IL-2 receptor (R) α−chain (*Il2ra*, CD25). IL-2 signalling induces further T cell activation, proliferation and differentiation (2). In addition, IL-2 signalling has key roles in immunological tolerance (2). This is partly mediated through CD25-expressing regulatory T cells (Treg), which suppress the activities of other T cells (3). Intriguingly, TCR signalling also induces the transient expression of FoxP3, the lineage-specific transcription factor of Treg (4), in any T cells in humans (5), and in mice in the presence of IL-2 and TGF-β (6). These suggest that FoxP3 can be actively induced as a negative feedback mechanism for the T cell activation process, especially in inflammatory conditions in tissues (7). Thus, the T cell activation processes may dynamically control Treg phenotype and function during immune response and homeostasis.

In fact, TCR signalling plays a critical role in Treg. Studies using TCR transgenic mice showed that Treg require TCR activation for *in vitro* suppression (8). The binding of Foxp3 protein to chromatin occurs mainly in the enhancer regions that have been opened by TCR signals (9). In fact, continuous TCR signals are required for Treg function, because the conditional deletion of the TCR-*α* chain in Treg abrogates the suppressive activity of Treg and eliminates their activated or effector-Treg (eTreg) phenotype (10, 11). It is, however, unclear how TCR signals contribute to the Treg-type transcriptional programme, and whether TCR signals are operating in all Treg cells or whether these are required only when Treg suppress the activity of other T cells.

Heterogeneity of Treg has been previously addressed through classifying Treg into subpopulations, according to the origin (thymic Treg, peripheral Treg, visceral adipose tissue Treg (12)), the transcription factor expression and ability to control inflammation (Th1-Treg (13) and Th2-Treg (14), and T follicular regulatory T cells (15)), and their activation status (activated Treg/eTreg, resting Treg, and memory-type Treg (16)). Among these Treg subpopulations, of interest is eTreg, which are activated and functionally mature Treg. Murine eTreg can be identified by memory/activation markers such as CD44, CD62L, and GITR (16, 17), and their differentiation is controlled by the transcription factors Blimp-1, IRF4 and Myb (18, 19). Human Treg can be classified into naïve Treg (CD25+CD45RA+Foxp3+) and eTreg (CD25+CD45RA-Foxp3+) (20). However, our recent computational study showed that classical gating approach is not effective for understanding multidimensional data, and that marker expression data may be rather effectively analysed by the computational clustering approaches that aim to understand the dynamics of marker expressions in Treg (21). Furthermore, the recent advancement of single cell technologies has opened the door to address the heterogeneity of Treg by their gene profiles at the single cell level.

When addressing the single cell level heterogeneity, it is critical to analyse activated effector T cells (Teff) and memory-like T cells (memory-phenotype T cells; Tmem) together with Treg. The surface phenotype of Tmem is CD44^high^CD45RB^low^CD25^−^ (22), which is similar to CD25^-^ Treg, apart from Foxp3 expression and suppressive activity (23, 24). In addition, Teff express CD25 and CTLA-4 (25), the latter of which is also known as a Treg marker (26). Tmem may include both antigen-experienced memory T cells (27) and self-reactive T cells (28). In fact, CD44^high^CD45RB^low^ Tmem do not develop in TCR transgenic mice with the *Rag* deficient background, indicating that they require agonistic TCR signals in the thymus (29). In addition, a study using a fate-mapping approach showed that a minority of Treg naturally lose Foxp3 expression and join the Tmem fraction (30). These suggest that, upon encountering cognate self-antigens, self-reactive T cells, which include Tmem and Treg, express and sustain Foxp3 expression as a negative feedback mechanism for strong TCR signals (7). Thus, Treg have a close relationship with Tmem and Teff. However, since most studies used naïve T cells (Tnaïve) as the control for Treg, many of known Treg-associated features may be in fact shared with Tmem and Teff.

Multidimensional analysis is an effective approach to address this problem, allowing to systematically investigate the relationships between more than two cell populations (e.g. based on transcriptional similarities) (31). The prototype methods include Principal Component Analysis (PCA), Correspondence Analysis (CA) (32) and Multidimensional Scaling (33). In the application to genomic data, these methods measure distances (i.e. similarities) between samples and/or genes using different metrics, and thereby visualise the relationships between samples and/or genes in a reduced dimension, typically either in two-dimensional (2D) or three-dimensional (3D) space, providing means to explore and investigate data (31). However, these multidimensional methods are often not sufficiently powerful for hypothesis-driven research, and our previous studies developed a transcriptome analysis method using a variant of CA, Canonical Correspondence Analysis (CCA) for microarray data (31) and RNA-seq data (34). In this approach, two transcriptome datasets are canonically analysed: the correlations between cell samples in one dataset (main dataset) and the immunological processes (explanatory variables) in another dataset (explanatory dataset) are analysed based on their correlations to individual genes. Briefly, CCA uses a linear regression to identify the interpretable part (constrained space) of main data by explanatory variables, and visualises similarities between genes, cells, and explanatory variables using a singular value decomposition (SVD) solution within the interpretable space (34). Thus, CCA enables to investigate and identify the unique features of each T cell population, visualising the relationships between T cell populations.

In this study, we investigate the multidimensional features of Treg in comparison to other CD4+ T cells including Teff, Tmem, and naïve T cells under normal or pathological conditions. Here we aim to identify the differential regulation of transcriptional modules for T cell activation and differentiation in these populations, and to reveal systems and molecular mechanisms behind the differential regulation. Furthermore, using our new single cell combinatorial CCA (SC4A) approach, we investigate the single-cell level heterogeneity of CD4+ T cells including Treg and effector T cells, identifying the differentially regulated gene modules and the bifurcation point for T cell fates.

## Materials and Methods

### Conventional CCA (Gene-oriented analysis)

CCA canonically analyses two independent microarray or RNA-seq datasets (34). Briefly, gene expression data of the standardised main dataset (**S**) is linearly regressed onto the explanatory variable(s) (**D**), which identifies the interpretable part of the main dataset (“Constrained data”, **S\***). When only one explanatory variable is used, the CA algorithm of CCA assigns numerical values to cell samples and genes so that the dispersion of samples is maximised (uncorrelated information components), providing a one-dimensional solution (34). The correlation analysis of explanatory data and gene scores in the CA solution generates a biplot value, which, in the one-dimensional solution, represents the explanatory variable score. In the one-dimensional CCA solution, the single biplot value can either be +1 or −1, indicating the direction (increasing/decreasing) of correlated genes in the explanatory variable against that in the main dataset. In order to use the one-dimensional solution as a scoring system, the CCA score (i.e. Axis 1 score) is multiplied by the single biplot value, which indicates positive or negative correlation to Axis 1, ensuring that cells and genes with high scores have high positive correlations to the explanatory variable. When two or more explanatory variables are used, the CA algorithm then performs SVD on **S**\*, creating new matrices (i.e. sample and gene score matrices). These scores are sorted into new uncorrelated axes α_k_, along which the entire set of scores generated by SVD is distributed. The first axis has the largest variation (*inertia*) and thereby explains the greatest amount of information extracted by the analysis. The map approach enables the comparison of two or more explanatory variables, while the regression process in CCA allows the analysis across two different experiments (34). Biplot values of the CCA result are shown by arrows on the CCA map. CCA provides a map that shows the correlations between samples of interest, explanatory variables, and genes. Highly correlated components are closely positioned on the map.

Note that the same genes must be used in both transcriptome dataset matrices. The main dataset is projected onto the explanatory variable dataset, thus the genes in common to both datasets comprise the interpretable part (intersect) of the main data. Mathematical operation implemented in the CCA algorithm produces immunological process (explanatory variable), gene and cell sample scores. The results are visualized as the 3-dimensional CCA solution on the CCA map (i.e. CCA triplot) that shows the relationships between cell subsets, genes and immunological processes. For example, in the application of CCA to population-level (bulk) data, transcriptome datasets of peripheral CD4+ T cells (including Treg, naïve, memory, draining LN and non-dLN and tissue effector CD4+ T cells) were processed by CCA using indicated explanatory variables (e.g. T cell activation) and the cross-level relationships between components at three different levels (immunological process, gene and cell) were analysed.

### Single cell data pre-processing and single cell CCA (single-cell oriented analysis)

RNA-seq expression data of GSE72056 was obtained from single-cell suspension of tumour cells with unknown activation and differentiation statuses (35). Genes with low variances and low maximal values were excluded. In order to identify CD4+ T cells, single cell data were filtered by the expression of *CD4* and *CD3E* to obtain only the *CD4*+*CD3E*+ single cells, and also by k-means clustering of PCA gene plot to exclude outlier cells (21) for subsequent analysis.

In the application of CCA to single cell data, importantly, the same single cells are used in both main data and the explanatory variables (i.e. selected genes). The main dataset is projected onto the explanatory variables, visualising the relationships between single cells, genes and explanatory variables, which represent major activation/differentiation processes in the dataset.

### Explanatory variables for conventional CCA

Explanatory variables for CCA were prepared as follows. Differentially-expressed genes were selected by a moderated t-test result using the Bioconductor package, *limma*. The top-ranked differentially expressed genes (according to their *p*-values) were used for making the explanatory variables. The T cell activation explanatory variable (*Tact*) was defined by the difference in gene expression between anti-CD3/CD28-stimulated (17 h) CD4^+^ T cells and untreated naïve CD4^+^ T cells from GSE42276 (36). Precisely, genes were selected by FDR < 0.01 and log2 fold change (> 1 or < −1) in the comparison of the gene expression profile of the activated and resting T cells. For the one-dimensional CCA of T cell populations (Figure 1B), the expression data of GSE15907 (37) was regressed onto gene values in *Tact* representing the change in gene expression following T cell activation, and CA was performed for the regressed data and the correlation analysis was done between the new axis and the explanatory variable. For the two-dimensional CCA of T cell populations (Figure 2A and 2B), the expression data of GSE15907 (37) was regressed onto *Tact*, in combination with *Foxp3* and *Runx1* explanatory variables representing the effects of *Foxp3* and *Runx1* expression on CD4+ T cells (GSE6939 (38)). *Foxp3* explanatory variable is the log2 fold change of *Foxp3*-transduced naïve CD4+ T cells and empty vector-transduced CD4+T cells. *Runx1* explanatory variable is the log2 fold change of *Runx1*-transduced naïve CD4+ T cells and empty vector-transduced CD4+T cells. Subsequently, CA was applied to the regressed data and the correlation analysis was performed between the new axes and the explanatory variables.

**Figure 1.**
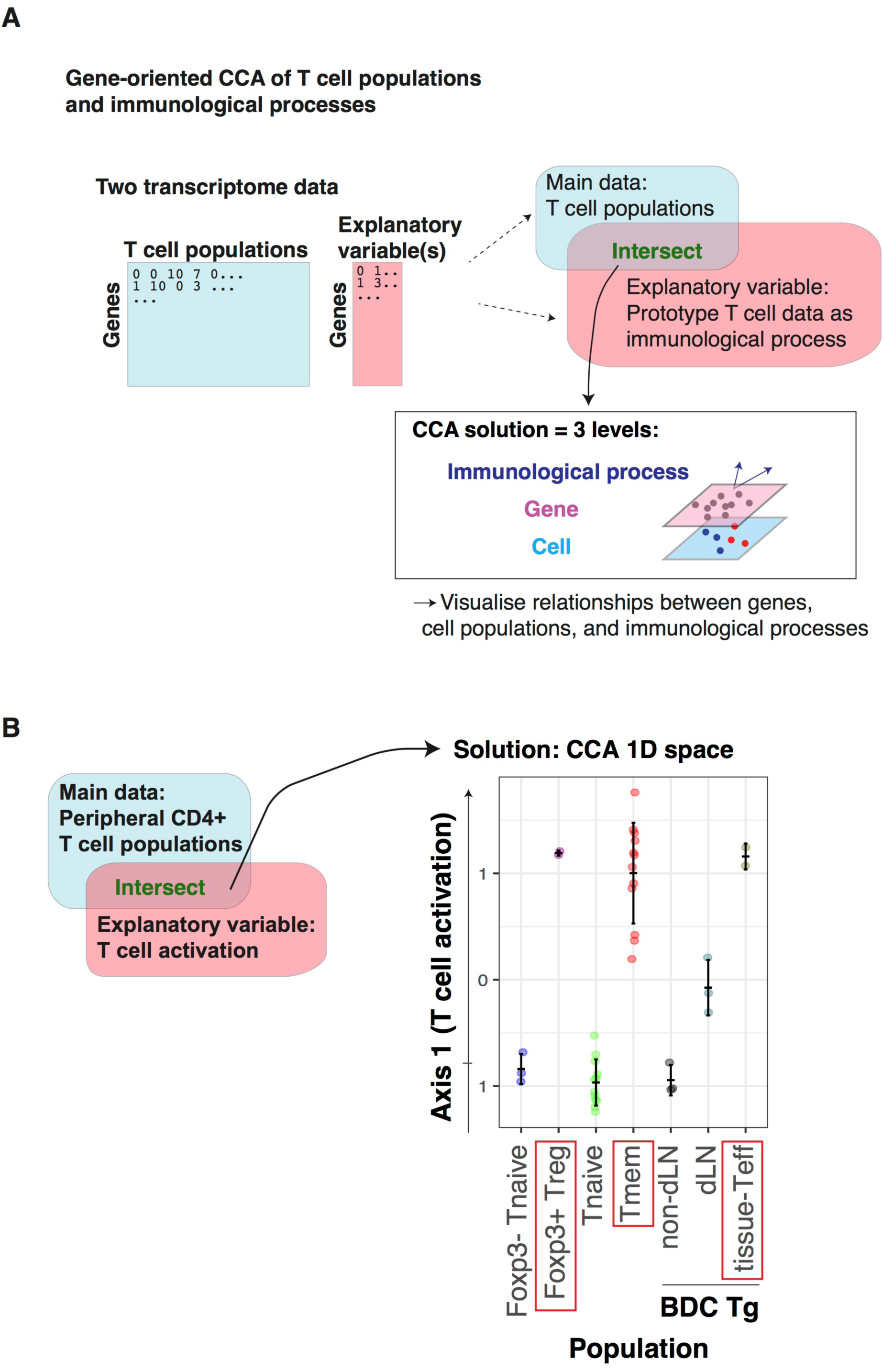
Identification of the activation signature in Treg and Tmem by CCA of T cell populations. The microarray dataset of peripheral CD4+ T cells, including naïve, effector and memory phenotype from various sites (GSE15907), was analysed using the T cell activation variable, which was obtained by the microarray dataset of conventional activated CD4+ T cells (GSE42276). (**A**) Schematic representation of CCA for the cross-level analysis of T cell populations (cells), immunological processes, and genes. (**B**) CCA was applied to the T cell population data using an explanatory variable for T cell activation, which was obtained as fold change between activated and resting conventional CD4+ T cells. The CCA solution is thus one-dimensional, and is used as “T cell activation score” (see Methods).

**Figure 2.**
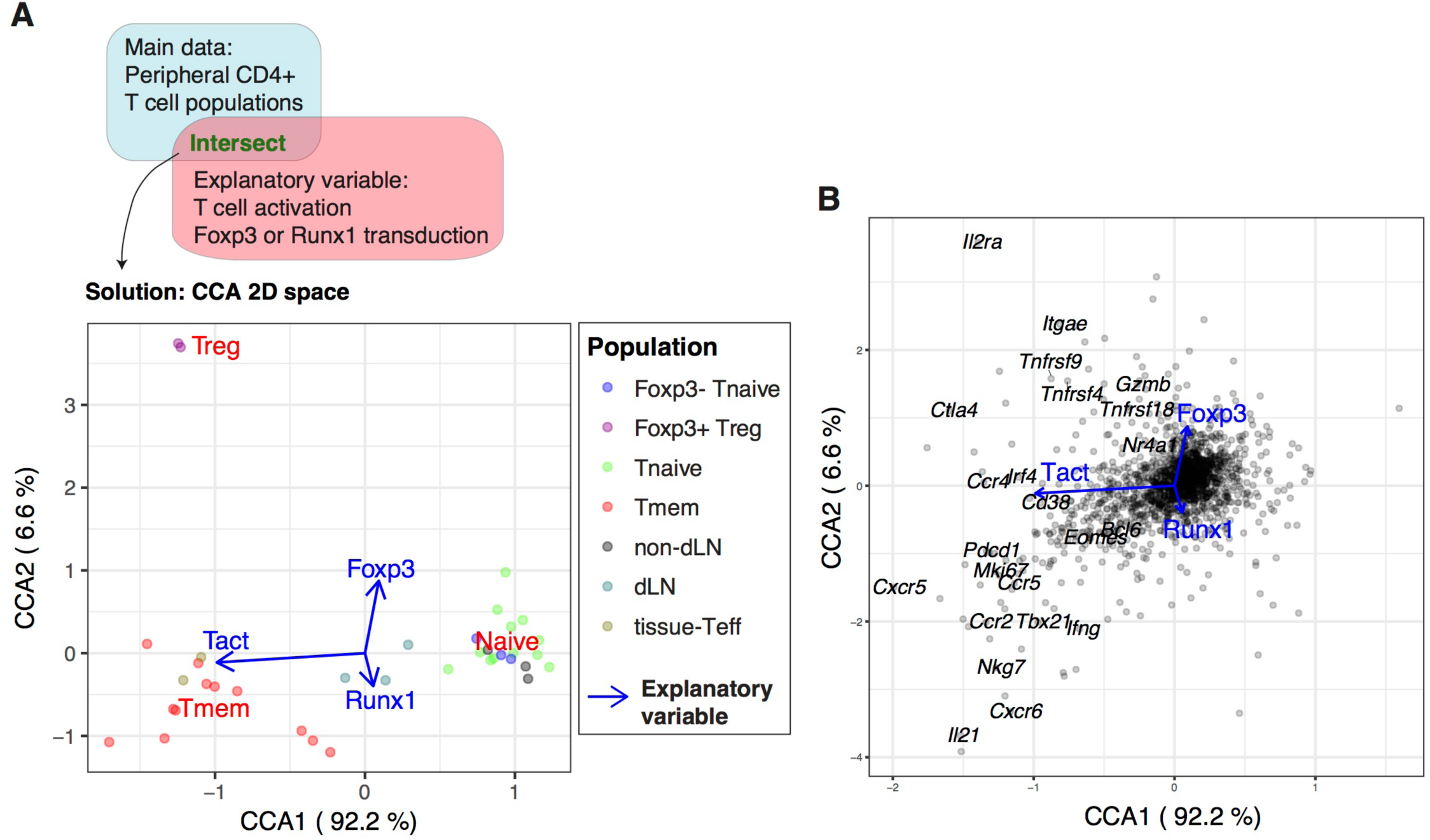
Identification of the Foxp3-independent activation signature in Treg by CCA of T cell populations. The microarray dataset of peripheral CD4+ T cells (GSE15907) was analysed using the T cell activation variable and the variables for retroviral *Foxp3* transduction and *Runx1* transduction as explanatory variables. (**A**) The CCA solution was visualised by a biplot where CD4+ T cell samples are shown by closed circles (see legend) and the explanatory variables are shown by blue arrows. Percentage indicates that of the variance accounted for by the inertia of the axes (i.e. the amount of information (eigenvalue) retained in each axis). (**B**) Gene biplot of the 2D CCA solution in (C) showing the relationships between genes (grey circles) and the explanatory variables (blue arrows). Selected key genes are annotated.

### SC4A

SC4A is a composite approach to understand non-annotated single cell data by identifying distinct populations of cells and the differentiation processes that are correlated with these populations. Since T cell population is usually identified by a single lineage specification factor, in the application of SC4A, such a factor will represent the cell population and their differentiation process (**Supplementary Figure 1A**). The advantage of this approach is that SC4A uses the gene expression data of a part of the dataset analysed, and thus the regression analysis of CCA becomes more efficient because of the absence of between-experimental variation, which is usually significant in cross-dataset analysis (34).

SC4A is performed by the following three steps: (1) identification of putative cell populations and candidate genes for explanatory variables by standard CCA; (2) combinatorial CCA to identify the top-ranked genes to be used as explanatory variables; and (3) the final CCA solution using the selected genes as explanatory variables.

#### 1. Preliminary analysis

The aim of the preliminary analysis is to identify the putative cell populations and candidate genes for explanatory variables, and standard CCA is a useful method to do this, because candidate genes can be identified by their correlation to each putative cell population. Considering that the final output is most effectively understood by visualisation using 2-dimensional (showing correlation between explanatory variable, samples and genes) or 3- dimensional (showing correlation between explanatory variable and samples/genes) plot, up to 4 cell populations will be identified, and up to 5- 10 genes for each population will be identified by their correlation to the population (**Supplementary Figure 1B**).

#### 2. Combinatorial CCA

Here SC4A aims to identify a set of genes that make the dispersion of cell populations maximum in the CCA solution. To achieve this, all the combinations of genes will be used as explanatory variables and tested for discriminating each two populations using CCA. During each combinatorial cycle, two genes are chosen from the total selected genes for all defined single-cell populations in the main dataset and tested for their correlations to one defined cell population vs all other T cells.

Correlation can be visualised as the degree angle measured between the explanatory variable (gene) and the centroid of the defined cell population. Out of the two genes, the gene with the smallest angle to the defined cell population is the most correlated. All selected genes are tested in this pairwise manner against all defined cell populations vs all other T cells to identify the gene that is most highly and uniquely correlated to each defined cell population. At each combinatorial CCA, the most correlated gene to each cell population is identified using vector multiplication (**Supplementary Figure 1C**). The top-ranked genes are determined by F1 score (the harmonic mean of precision and sensitivity) and the correlation to the population of interest. When the top-ranked gene is different between F1 score and the correlation, the most immunologically-meaningful gene can be chosen.

#### 3. SC4A

Finally, CCA is performed using the genes that are selected by the combinatorial CCA to be used as explanatory variables. Thus, the single cell dataset will be explained by the expression of the set of chosen genes, each of which uniquely correlates with a cell population and represents the differentiation process of the cell population (**Supplementary Figure 1D**).

Algebraically, SC4A is defined as follows. Single cell RNA-seq data **X** ∈ **R***^p^ ^×^ ^m^* is the measurement of *m* genes from *p* single cells. The *j*-th column *x_j_* = (x_1j_ x_2j_ … x_pj_)^T^ is the expression data of the j-th gene from *p* single cells, where T indicates transposed vector. In SC4A, by choosing a set of *k* genes for explanatory variable, **X’** ∈ **R***^p^ ^×^ ^(m-k)^* will be analysed by CCA using **Z** ∈ **R***^p^ ^×^ ^k^* as explanatory variables. As in the algorithm for CCA, **X’** is standardised by column sums (***c***) and row sums (***r***), i.e. **S** = **D_r_**^-1/2^ (*1/n* **X**– **rc**T) **D_c_**^-1/2^, where *n* is the grand total of gene expression data, **D_r_** and **D_c_**are the diagonal matrices of **r** and **c**, respectively. Meanwhile, **Z** is scaled and standardised (i.e. mean = 0 and variance = 1). **S** is linearly regressed onto **Z** by the projection matrix **Q** = **D_r_**^1/2^ **Z** (**Z**^T^ **D_r_ Z**)^−1^ **Z**^T^ **D_r_**^1/2^, and the constrained space **S\*** = **Q S**. Next, CCA performs singular value decomposition (SVD) of **S\*** = **U D_α_ VT**, where **UT U** = **VT V** = **I**, and **D_α_** is the diagonal matrix of singular values in descending order (α1 ≥ α2 ≥ …). Thus, SVD analyses the constrained space and provides new axes where the dispersion of samples and that of genes are maximised in the first axes. Gene scores are defined as **D_r_^-1/2^ U D_α_** or **D_r_^-1/2^ U**, and sample scores for single cells are defined by weighted average scores (WA scores) **S V D_α_**, or **S V**.

#### 4. Choice of explanatory variables by SC4A

SC4A aims to identify a set of genes that make the dispersion of cell populations maximum in the CCA solution. To achieve this, all the combinations of genes will be used as explanatory variables and tested for discriminating each two populations using CCA. During each combinatorial cycle, two genes are chosen from the total selected genes for all defined single-cell populations in the main dataset and tested for their correlations to one defined cell population vs all other T cells. In the analysis of Figure 8, the following two cell populations were analysed by the combinatorial CCA: (1) Activated T cells vs Resting T cells; (2) FOXP3+ cells vs FOXP3- cells; (3) BCL6+ cells (as Tfh-like T cells) vs BCL6- cells. The most correlated gene to each population (Activated T cells, Resting T cells, FOXP3+ cells, or BCL6+ cells) was identified, and these 4 genes were used as explanatory variables in the final output of SC4A in Figure 8.

### Data pre-processing and other statistical methods

All microarray datasets were downloaded from GEO site, and normalized, where appropriate using the Bioconductor package *Affy*. Data were arranged into an expression matrix where each row corresponds with gene expression for each gene and each column corresponds with cell phonotype (sample). Data were log2-transformed and values above log2(10) were used for analysis. Differentially expressed genes (DEG) the TCR KO dataset and the aTreg dataset were identified by a moderated t-statistics. DEG for activated CD44hi and resting CD44lo Treg were combined. The CRAN package *vegan* was used for the computation of CCA. Gene scores used the *wa* scores of the CCA output by vegan. The Bioconductor package *limma* was used to perform a moderated t-test. RNA-seq data were preprocessed, normalised, and log-transformed using standard techniques (34).

Heatmaps were generated the CRAN package *gplots*. Venn diagram was generated using the R code, overLapper.R, which was downloaded from the Girke lab at Institute for Integrative Genome Biology (http://faculty.ucr.edu/~tgirke/Documents/R_BioCond/My_R_Scripts/overLapper.R). Gene lists were compared for enriched pathways in the REACTOME pathway database using the Bioconductor packages *ReactomePA* and *clusterProfiler*. Violin plots shows kernel density plots (outside) and the median and interquartile range (inside) of the original gene expression data, and were generated by the Bioconductor package *ggplot2*. The lineage curve was constructed by clustering SC4A/CCA sample scores using an expectation–maximization (EM) algorithm (39), and the nodes of these clusters were identified by constructing a minimum spanning tree using the Bioconductor package *Slingshot* (40).

## Results

### Identification of the Foxp3-independent activation signature in Treg and memory-phenotype T cells

Firstly, we investigated how T cell activation-related genes are differentially regulated in resting Treg and other CD4+ T cell populations including Tmem and Teff. To address this multidimensional problem, we applied CCA to the microarray dataset of various CD4^+^ T cells using the explanatory variable for the T cell activation process, which was obtained from the microarray dataset that analysed resting and activated conventional T cells (“T cell subset data” and “T cell activation data” in **Table 1**). Thus, we aimed to visualise the cross-level relationships between genes, the T cell populations, and the T cell activation process (**Figure 1A**). Using the single explanatory variable, the T cell activation process, the solution of CCA is one-dimensional and the cell sample scores of CCA (represented by Axis 1) provides “T cell activation score” (see Methods), indicating the level of activation in each cell population relative to the prototype signature of T cell activation, as defined by the explanatory variable *Tact*. All the naïve T cell populations had low Axis 1 values (i.e. Foxp3- T naïve cells (Tnaive); Tnaive, and non-draining lymph node (dLN) T cells from BDC TCR transgenic (Tg) mice, which develop type I diabetes). In contrast, Foxp3^+^ Treg, Tmem, and tissue-infiltrating Teff in the pancreas from BDC Tg (i.e. with inflammation in the islets) had high scores (**Figure 1B**). These results indicate that Treg are as “activated” as Tmem and tissue-infiltrating activated Teff at the transcriptomic level by CCA.

**Table 1.**
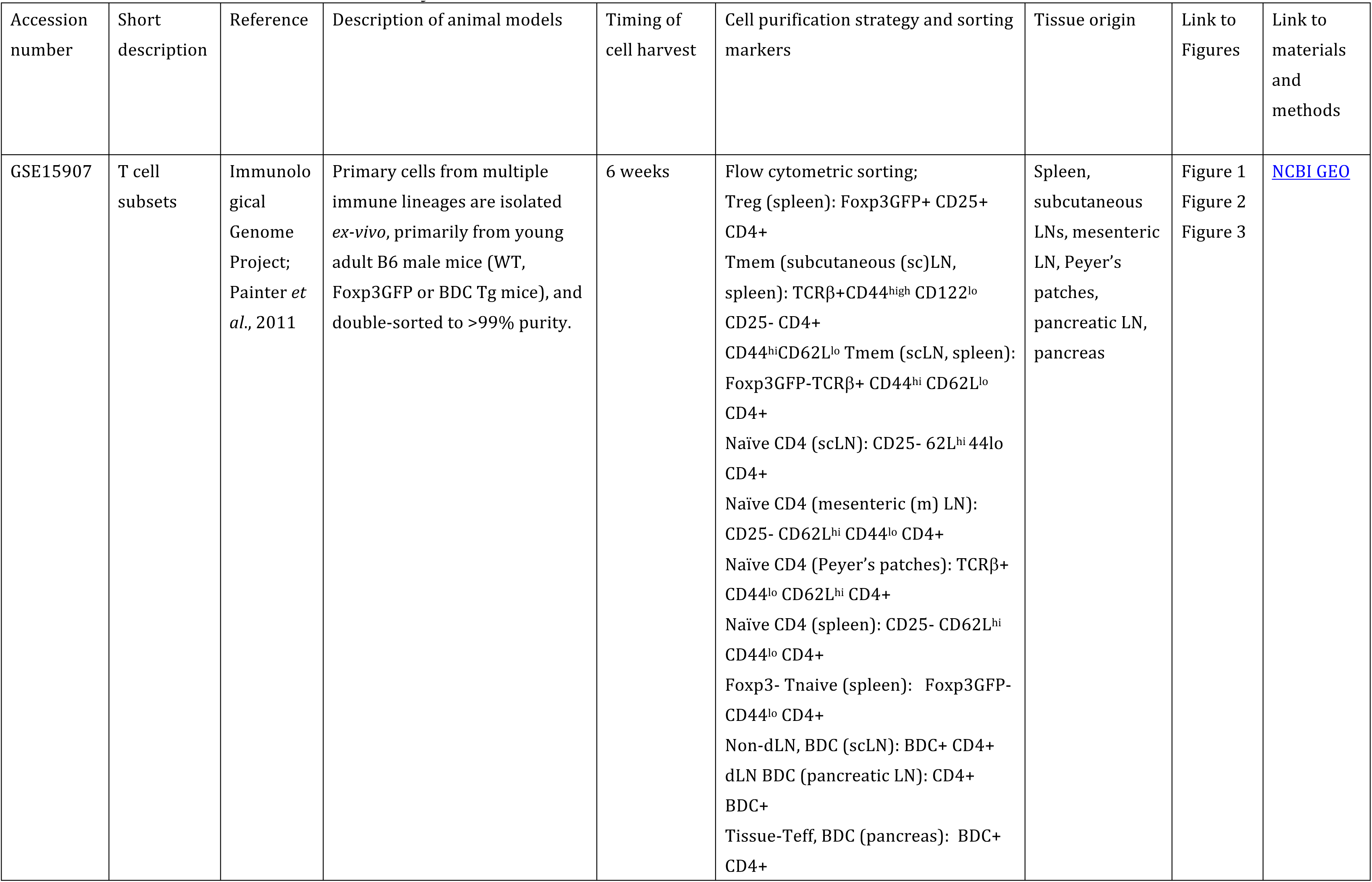
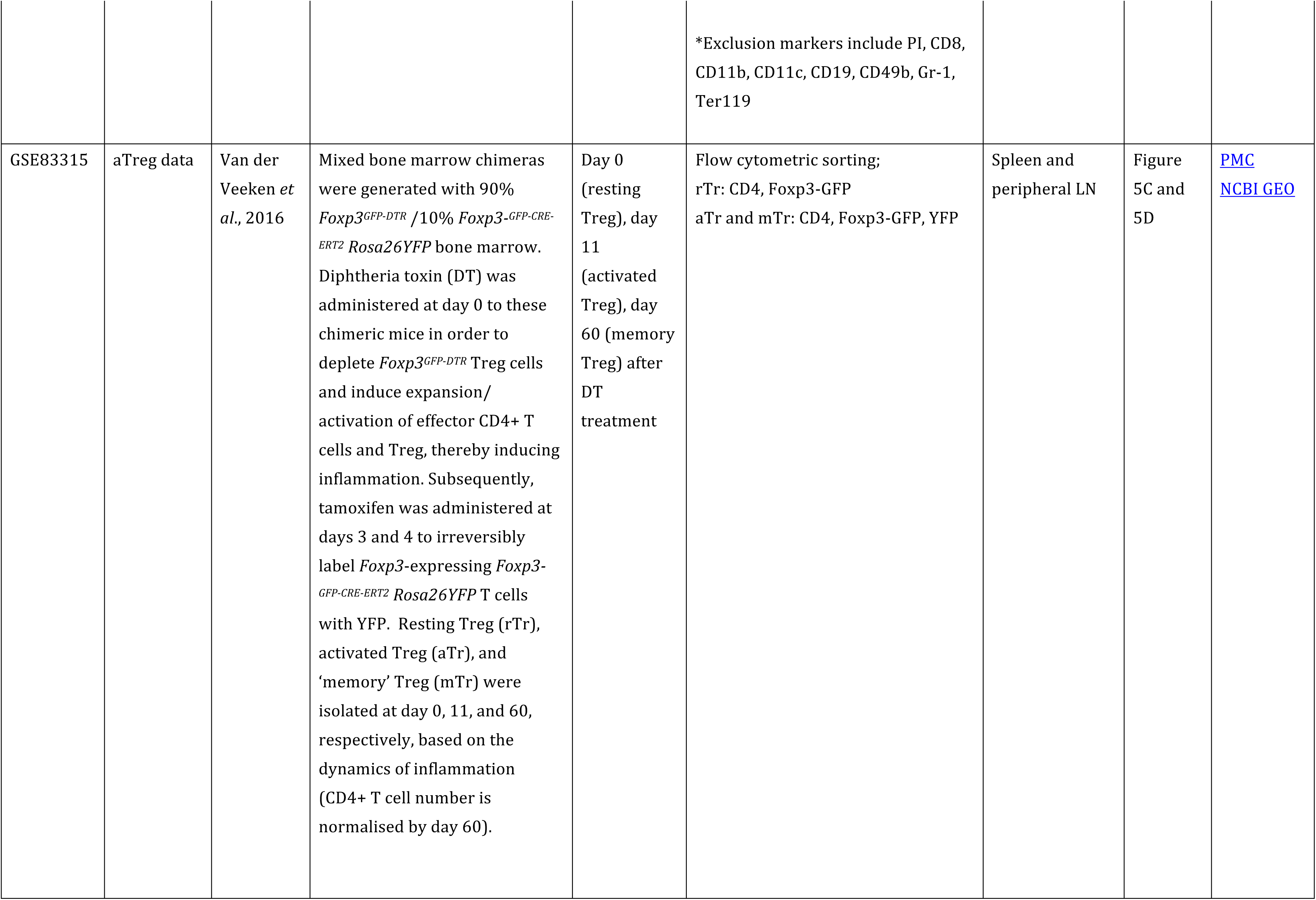
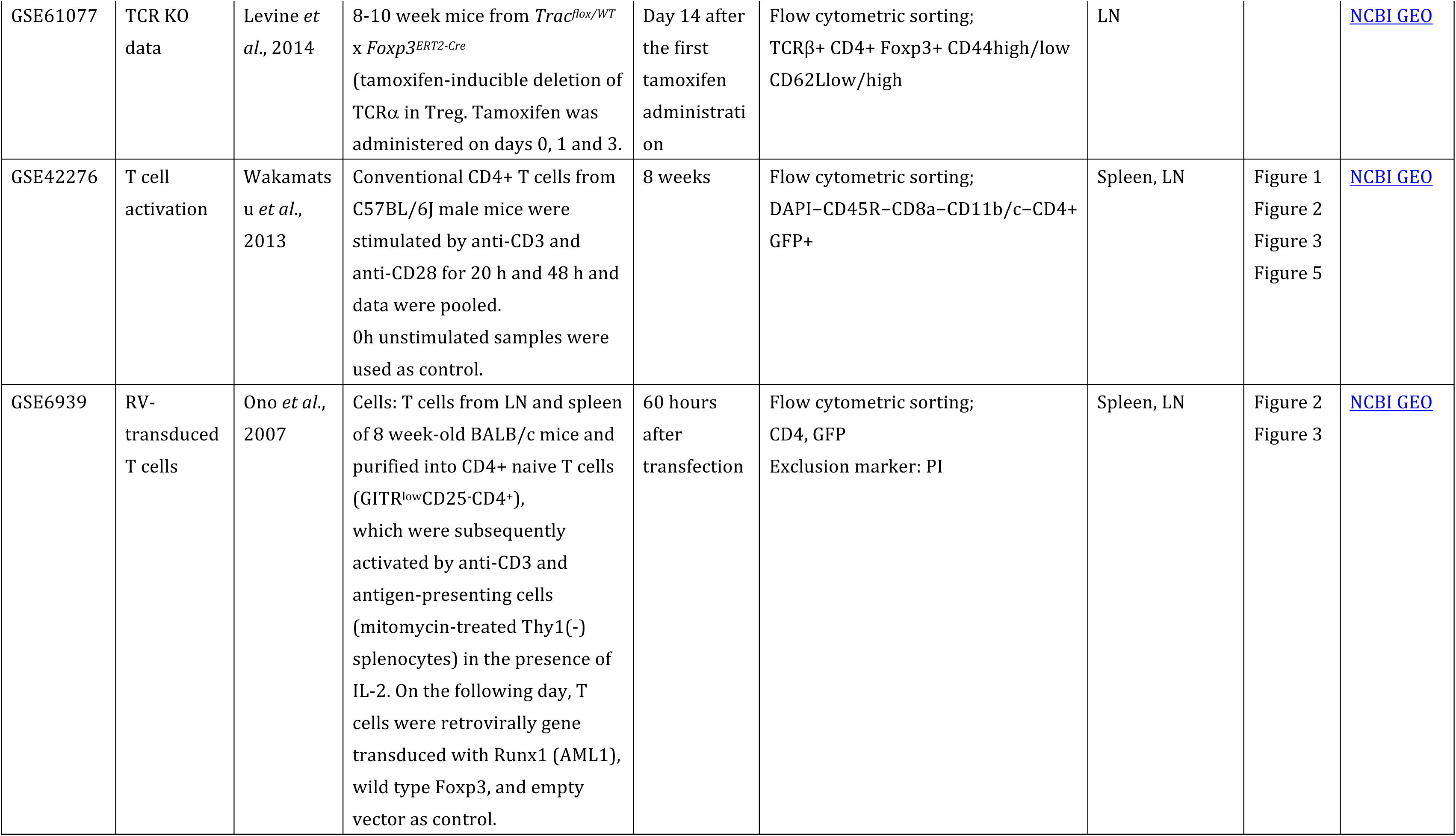
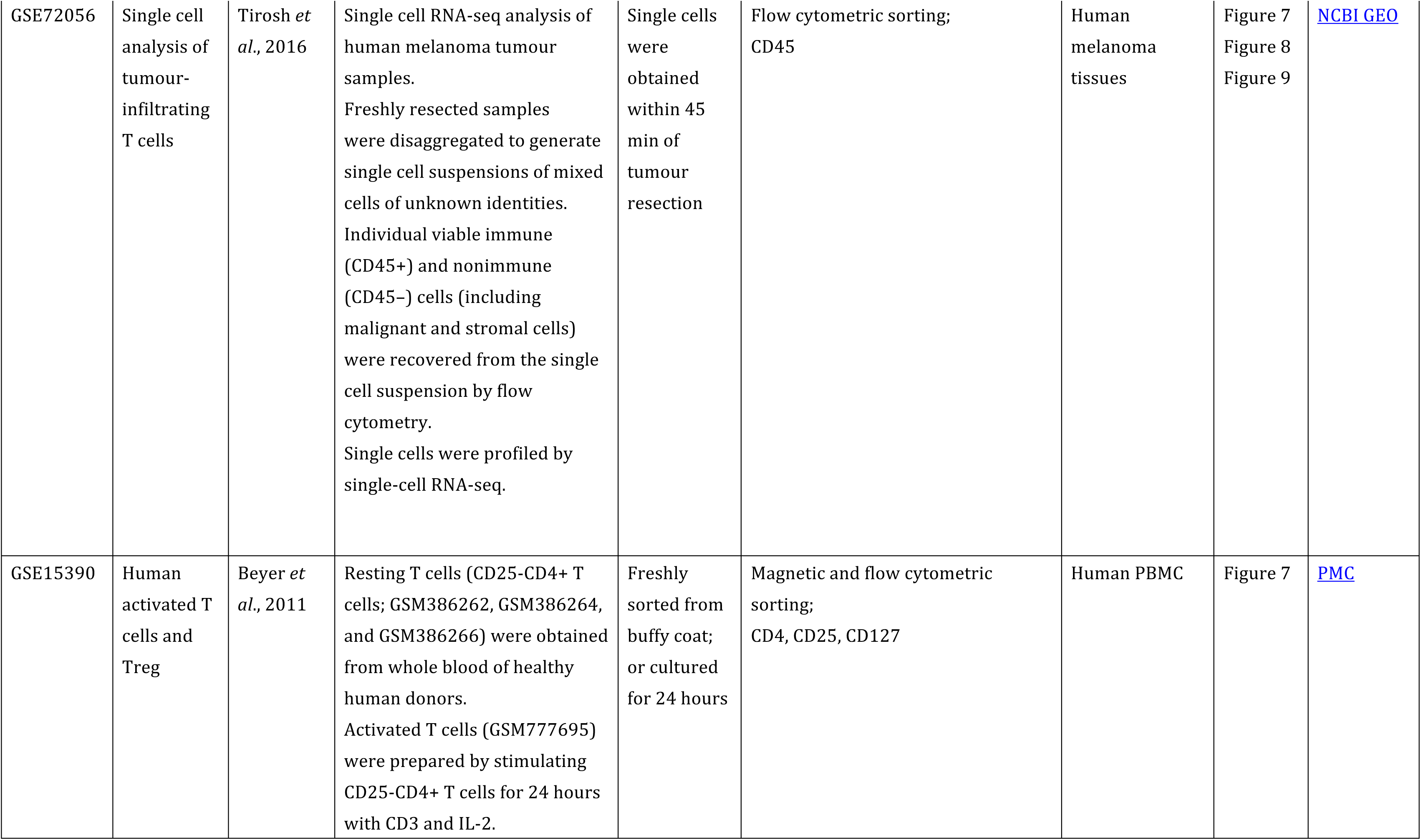
Datasets used in this study

Next, we addressed whether the highly “activated” status of Treg is dependent on Foxp3. Since Foxp3 suppresses Runx1-mediated transcriptional activities (38), we investigated the same T cell population dataset using the following three explanatory variables: T cell activation (Tact), retroviral *Foxp3* transduction (Foxp3) and *Runx1* transduction (Runx1) (see Methods). The CCA solution was 3-dimensional, while the first two axes explained the majority of variance (98.8%, **Figure 2A**). As expected, Tmem, tissue-infiltrating Teff and Treg had low negative values and showed high correlations to T cell activation (Tact) in Axis 1, whereas only Treg had high correlations with the Foxp3 variable in Axis 2, while Tmem and Teff were correlated with the Runx1 variable in Axis 2 (**Figure 2A**). By analysing the gene space of the CCA solution, genes in the lower left quadrant (i.e. negative in both Axes 1 and 2) were enriched with the genes that are involved in T cell activation, effector functions, and T follicular helper cells (Tfh), including *Cxcr5*, *Pdcd1(*PD-1) *Il21, Ifng, Tbx21* (T-bet)*, Mki67* (Ki-67) (**Figure 2B**). On the other hand, genes in the upper left quadrant (i.e. negative in Axis 1 and positive in Axis 2) were enriched with Treg-associated genes including *Ctla4*, *Il2ra* (CD25), *Itgae* (CD103), *Tnfrsf9* (4-1BB) and *Tnfrsf4* (OX40) (**Figure 2B**). These results indicate that a set of activation genes are operating in all the three non-naïve T cell populations (i.e. Treg, Teff and Tmem), while some of them are more specific to Treg.

### The Treg transcriptome is characterized by the repression of a part of the activation genes for Tmem

Next, we determine the modules of genes that are differentially regulated between Treg and Tmem, in order to understand the multidimensional identity of Treg and Tmem transcriptomes (i.e. how these populations can be defined in comparison to all relevant populations). Specifically, we asked if the Axis 2 captured the differential transcriptional regulations between Tmem and Treg. Importantly, Axis 2 represents Foxp3-driven and Runx1-driven transcriptional effects, which are correlated with Treg and Tmem/Teff, respectively (**Figure 3A**). This suggests that Axis 2 provides a ‘scoring system’ for regulatory vs effector functions. Thus, the genes in Axis 1-low (precisely, genes above 25 percentile for positive correlations with Tact) were identified as *Tact genes*. These genes were subsequently classified into Axis 2-positive (i.e. positive correlations with Foxp3 and Treg) [designated as “*Tact-Foxp3 genes*”; top left quadrant of CCA gene space in Figure 1D] and Axis 2-negative genes (i.e. positive correlations with Runx1 and Tmem/Teff) [designated as “*Tact-Runx1 genes*”; bottom left quadrant of CCA gene space in Figure 1D] (**Figure 3A**). Tact-Runx1 genes contain genes linked to T cell activation (e.g. *Mki67*), effector functions (e.g. *Tbx21*), and Tfh (e.g. *Bcl6*, *Pdcd1*), while Tact-Foxp3 genes contain “Treg markers” such as *Il2ra* (CD25) and *Tnfrsf18* (GITR) (**Figure 2B**).

**Figure 3.**
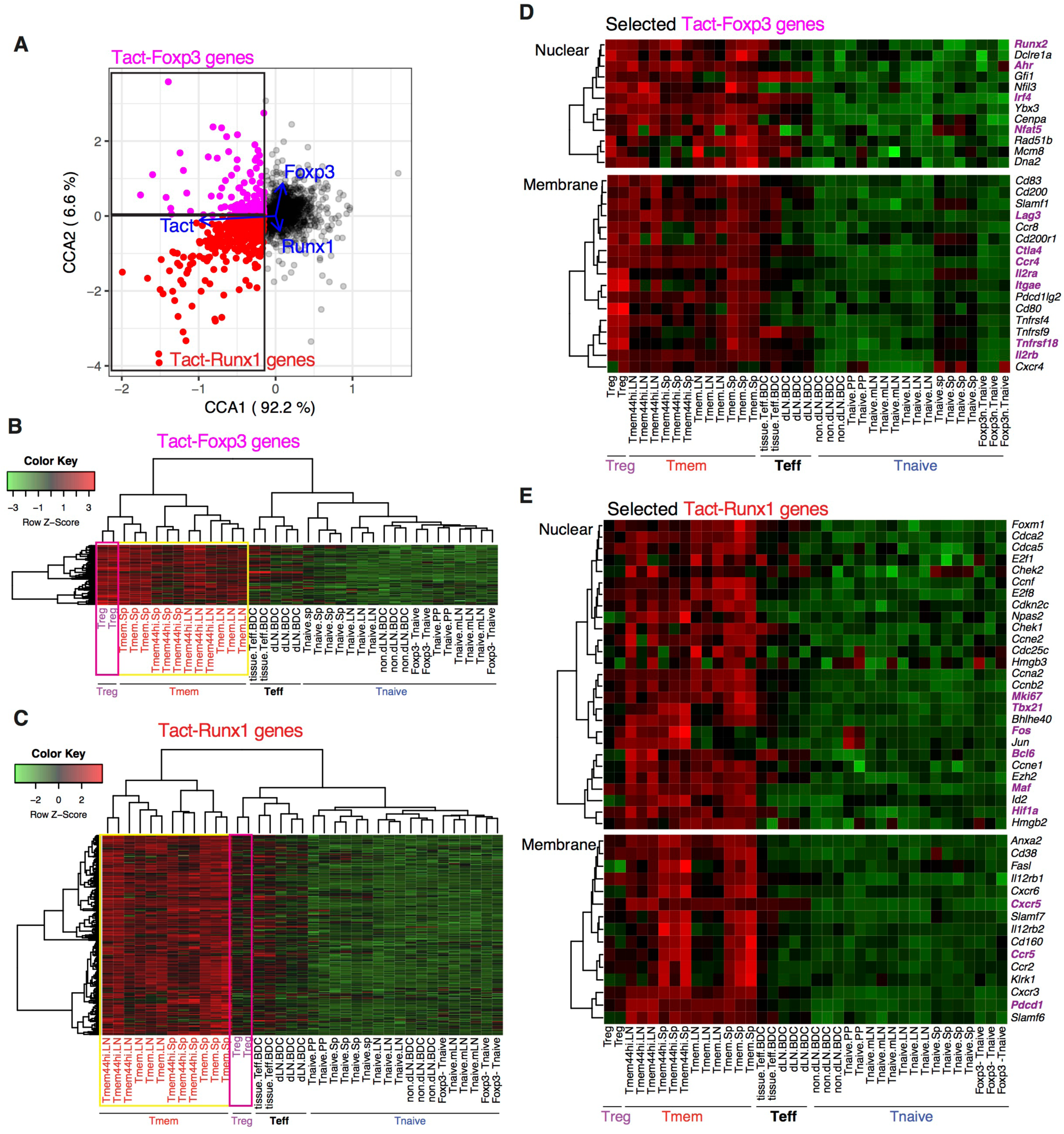
Differential regulations of transcriptional modules for activation in Treg and Tmem by Foxp3 and Runx1. (**A**) Definition of Tact-Foxp3 genes and Tact-Runx1 genes. In the gene plot of the CCA solution in Figure 2B, Axis 1-low genes (25 percentile low) were designated as *activation genes*, which were further classified into Tact-Foxp3 genes and Tact-Runx1 genes by Axis 2, which have high correlations to Treg and Tmem samples, respectively, in the CCA cell space (Figure 2A). (**B**) Heatmap analysis of all the Tact-Foxp3 genes. (**C**) Heatmap analysis of all the Tact-Runx1 genes. (**D**) Heatmap analysis of selected Tact-Foxp3 genes. (**E**) Heatmap analysis of selected Tact-Runx1 genes.

Intriguingly, heatmap analysis showed that both Treg and Tmem expressed Tact-Foxp3 genes at high levels, compared to naïve and effector T cells (**Figure 3B**). On the other hand, Tact-Runx1 genes were selectively downregulated in Treg, while their expressions were sustained in Tmem (**Figure 3C**). In other words, the repression of Tact-Runx1 genes was the major feature of Treg in comparison to Tmem, and Tact-Foxp3 genes are the activation genes, the expression of which is induced by T cell activation in both Treg and Tmem, and is sustained or enhanced even in the presence of Foxp3. Interestingly, comparable selective downregulation of Tact-Runx1 genes was observed in Teff as well (**Figure 3C**). This suggests that the set of activation genes operating in Teff is different from the ones in Tmem, and that Tmem and Treg share more activation genes than Treg-Teff and Tmem-Teff (**Figure 3B** **and 3C**). These results collectively indicate that the Treg-ness is composed of the induction of the Treg-Tmem shared activation genes (i.e. Tact-Foxp3 genes) and the Foxp3-mediated repression of Tmem-specific genes (i.e. Tact-Runx1 genes), defining the multidimensional identity of Treg.

While the overall activation levels of Treg and Tmem are similar to the ones of the tissue-infiltrating Teff at transcriptional level (Figure 1B), when explained by the prototype signature of activation in CD4+ T cells (i.e. the explanatory variable Tact), the compositions of the activation genes are different between Treg, Tmem and Teff (as captured by Figure 3B and 3C). Importantly, many of these activation genes are shared between Treg and Tmem, but not with Teff. The closer similarity between resting Treg and Tmem, compared to Teff, is not surprising, considering that both resting Treg and Tmem are at the resting status, while Teff are more recently activated and executing effector functions. In addition, the distinct features of Teff may also include their capacity of tissue infiltration and the effects of the microenvironment. These features were not captured by standard t-test analysis (**Supplementary Fig 1**).

Tact-Foxp3 genes included the transcription factors *Nfat5*, *Runx2*, and *Ahr,* which were expressed by most of Tmem cells as well (**Figure 3D**). The Treg-associated markers, *Il2ra* (CD25), *Itgae* (CD103), and *Tnfrsf18* (GITR) were expressed not only by Treg but also by Tmem at moderate to high levels. Notably, the expression of *Ctla4, Ccr4, and Lag3* was high in Treg and Tmem cells, but it was repressed in Teff (**Figure 3D**). This suggests that Treg and Tmem are in later stages of T cell activation, when the expression of CTLA-4 is induced as a negative feedback mechanism (41), while it is not induced in tissue-infiltrating Teff, presumably because they are more recently activated and actively proliferating.

Tact-Runx1 genes included many cell cycle-related genes (e.g. *Ccna1*, *Cdca2*, and *Chek2*), suggesting that these cells are in cell cycle and proliferating (**Figure 3E**). The higher expression of *Mki67* and *Fos* suggests that these Tmem cells had been activated by TCR signals *in vivo* before the analysis. Tact-Runx1 genes also included the transcription factors *Tbx21*, *Maf*, *Hif1a*, and *Bcl6*, which have roles in Th1, Th2, Th17, and Tfh differentiation, respectively (42–44). In accordance with this, the Tfh markers *Cxcr5* and *Pdcd1* were specifically expressed by Tmem, suggesting that Tmem are heterogenous populations and composed of these Th and Tfh cells. These results are compatible with the model that Treg and Tmem constitute the self-reactive T cell population that have constitutive activation status (7), and that the major function of Foxp3 is to modify the constitutive activation processes by repressing a part of the activation gene modules (i.e. Tact-Runx1 genes) (**Figure 4**).

**Figure 4.**
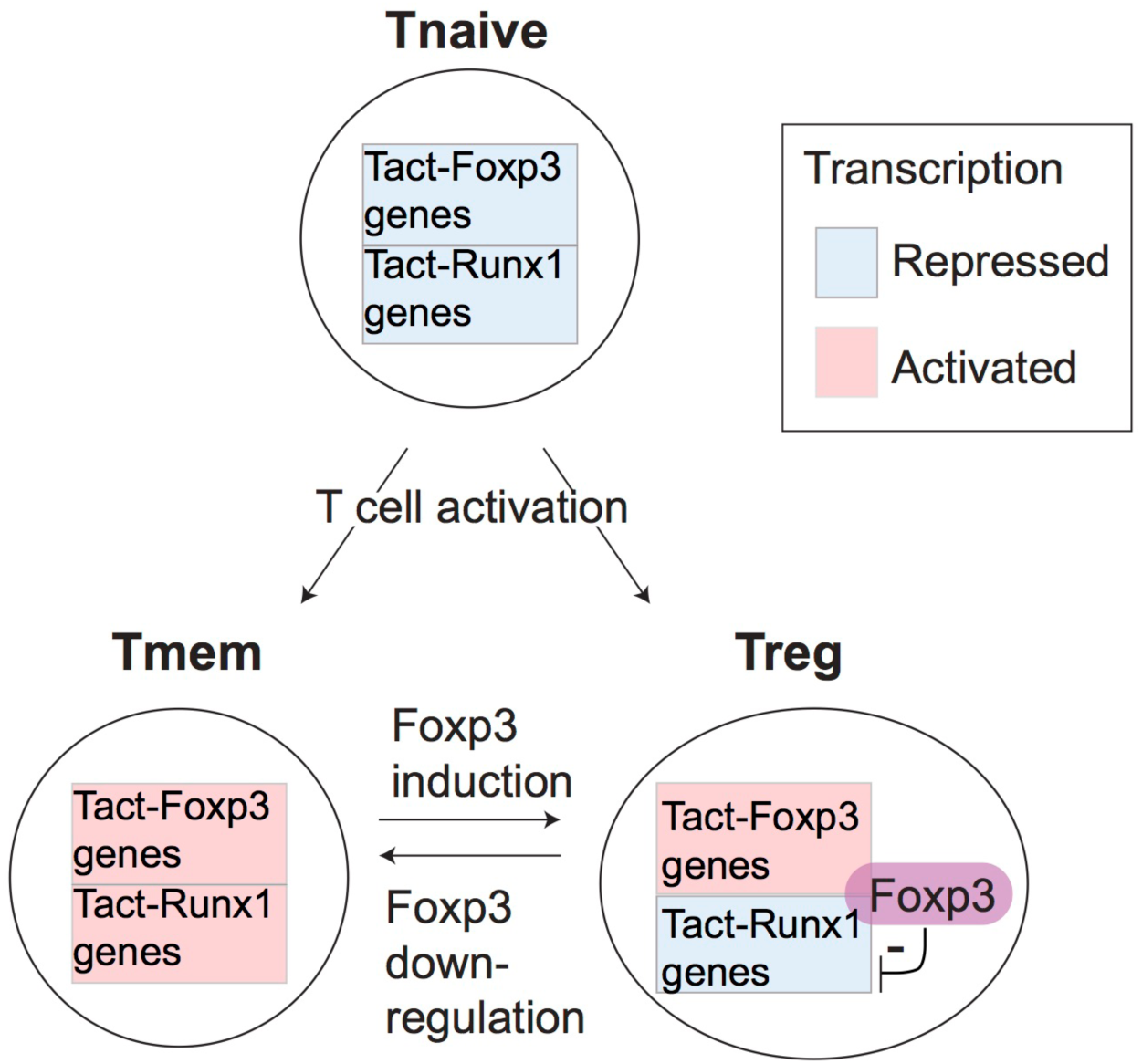
A model for the differential regulation of activation genes in Treg and Tmem. The proposed differential regulations of TCR signal downstream genes in Treg and Tmem. Since both naturally-arising Treg and Tmem are self-reactive T cells, they may frequently receive tonic TCR signals by recognising their cognate antigens in the periphery. This results in the full activation of both the Tact-Foxp3 and Tact-Runx1 gene modules in Tmem. However, in Treg, Foxp3 represses Tact-Runx1 genes and sustains the expression of Tact-Foxp3 genes, producing the characteristic Treg transcriptome.

### The activated status of Treg is TCR signal dependent

We next asked whether the constitutively “activated” status of Treg is dependent on TCR signals. We applied CCA to the microarray data of CD44^hi^CD62L^lo^ activated Treg (CD44^hi^ activated Treg) and CD44^lo^CD62L^hi^ naïve-like Treg (CD44^lo^ naïve Treg) from inducible *Tcra* KO or WT (TCR KO data, **Table 1**, **Figure 5A**) using the T cell activation variable as explanatory variable. The CCA result showed that CD44^hi^ activated Treg from WT mice only showed high activation scores, compared with all the other groups. Interestingly, *TCRa* KO CD44^lo^ naïve-like Treg showed the lowest scores, and were lower than WT CD44^lo^ naïve-like Treg (**Figure 5B**). These results indicate that TCR signaling is required for the constitutive activation status of Treg, especially CD44^hi^ activated Treg, and suggest that these activated Treg are more enriched with the cells that received TCR signals recently, compared to CD44^lo^ naïve-like Treg.

**Figure 5.**
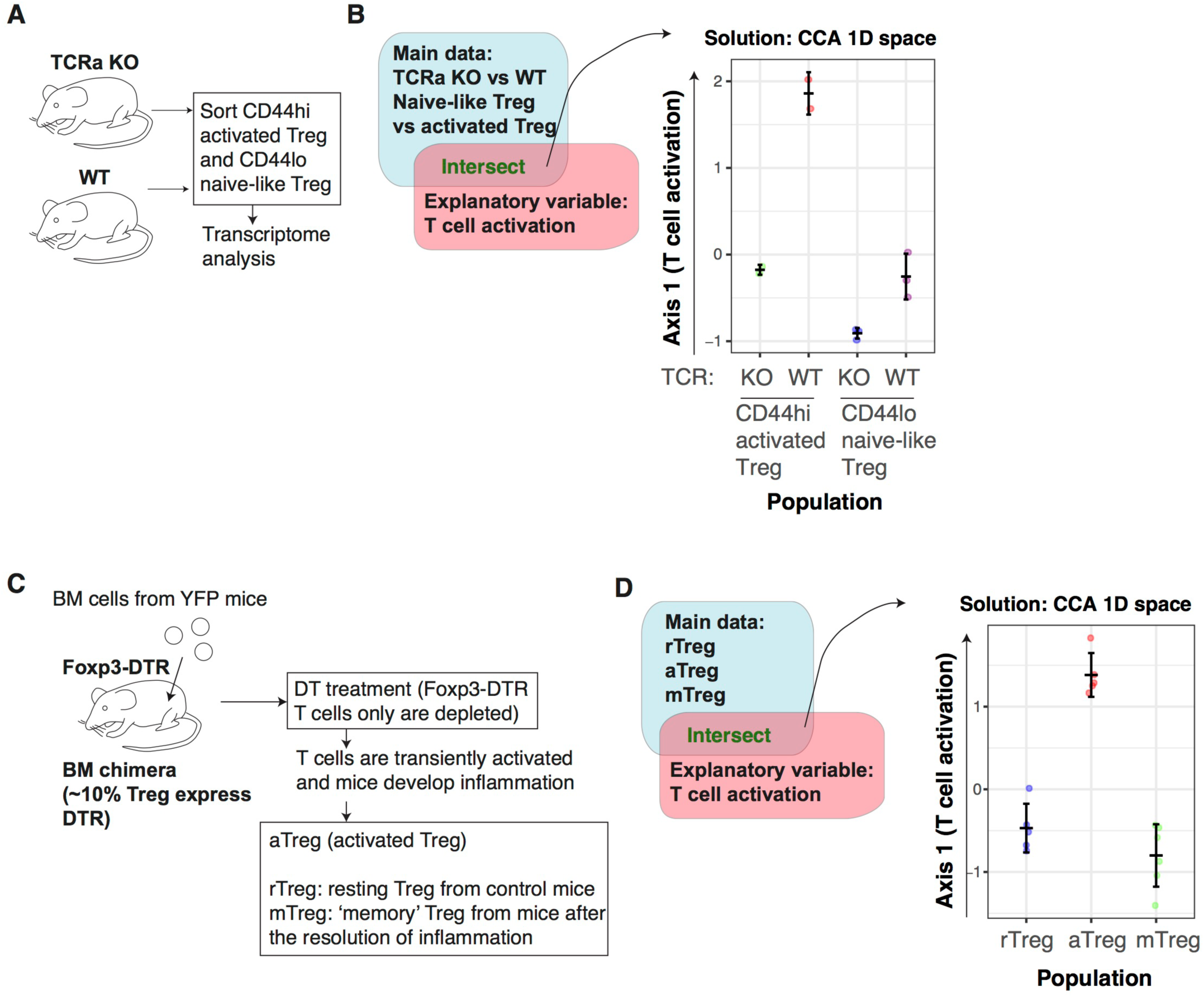
The activation signature of Treg is dependent on TCR signalling. (**A**) The experimental design for the TCR dataset. CD44hi activated Treg and CD44lo naïve-like Treg were obtained from TCRa KO or WT mice and analysed by transcriptome analysis. (**B**) CCA was applied to the transcriptome data of CD44^lo^CD62^hi^ naïve-like and CD44^hi^CD62^lo^ activated Treg cell populations from inducible *TCRa* KO or WT (from the *TCR KO* data, GSE61077), using the T cell activation variable as the explanatory variable. This produces a 1D CCA solution, and the sample score was plotted (representing “T cell activation score”). (**C**) The experimental design for the activated Treg dataset. Bone marrow (BM) cells were obtained from *Foxp3^GFPCreERT2^:Rosa26YFP* mice (YFP mice), and transferred into *Foxp3^GFP^ ^DTR^* mice (Foxp3-DTR mice), in order to make BM chimera, in which ∼10% of Treg expressed DTR. Subsequently, DT was administered to these BM chimera, which depleted Foxp3-DTR cells but not donor cells. This treatment induced a transient activation of T cells and inflammation *in vivo*. Activated Treg (aTreg) were obtained from these mice with inflammation, while resting Treg (rTreg) were from control mice, and memory Treg (mTreg) were from the mice after the resolution of inflammation. (**D**) 1D CCA sample score plot of transcriptomic data of resting Treg (rTreg), *in vivo* activated Treg (aTreg) and memory Treg (mTreg) from the *aTreg* data (GSE83315), with T cell activation signature as explanatory variable.

In order to further address whether the TCR signal-dependent activation signature of Treg is constitutively maintained or specifically induced by *in vivo* activation events (presumably as tonic TCR signals (7)), we analysed the RNA-seq dataset of *in vivo* activated Treg (Ref. (16), **Table 1**). The dataset was generated by depleting a part of Treg by Diphtheria toxin (DT) using bone marrow chimera of *Foxp3^GFPCreERT2^:Rosa26YFP* and *Foxp3^GFP^ ^DTR^* (16). The DT treatment depletes DT receptor (DTR)-expressing Treg from *Foxp3^GFP^ ^DTR^*, and thus induces a transient inflammation through the reduction of Treg. *Foxp3^GFPCreERT2^* allows to label Foxp3-expressing cells by YFP at the moment of tamoxifen administration. Van der Veeken *et al* thus analysed resting Treg from untreated mice (rTreg), activated Treg from mice with recent depletion (11 days before the analysis) in an inflammatory condition (aTreg), and “memory” Treg (mTreg) from mice with a distant depletion (60 days before the analysis) (**Figure 5C**). As expected, the CCA analysis using the T cell activation variable showed that aTreg had higher activation scores than both rTreg and mTreg (**Figure 5D**). This indicates that the activation mechanisms are more actively operating in activated Treg in an inflammatory environment.

In order to further dissect the activation signature of Treg, we obtained the lists of differentially expressed genes (DEG) between WT Treg vs *Tcra* KO Treg (designated as *TCR-dependent genes*), and between aTreg and rTreg (designated as *aTreg-specific genes*, see Methods). Interestingly, 94/286 genes of Tact-Runx1 genes (Tmem-specific activation genes, repressed in resting Treg) are also used during the activation of Treg (**Figure 6A**), while only 8/119 of Tact-Foxp3 genes (used by Tmem and resting Treg) are induced during the activation of Treg (**Figure 6B**). This indicates that the activation of Treg does not enhance the genes that are used in resting Treg, but induces the expression of the Tmem-specific genes that are suppressed in resting Treg. On the other hand, 51/286 of Tact-Runx1 and 19/119 of Tact-Foxp3 genes are regulated by TCR signalling (**Figure 6A and 6B**), suggesting that the activation status of resting Treg and Tmem may be sustained by TCR signals. Pathway analysis showed that Tact-Runx1 and aTreg-specific genes were enriched for cell-cycle related pathways. In contrast, Tact-Foxp3 genes were enriched for pathways related to signal transduction only (**Figure 6C**). Collectively, the results above suggest that resting Treg are maintained by TCR and cytokine signalling, and that the activation of Treg induces the transcriptional activities of Tact-Runx1 genes, which promote proliferation and cell division.

**Figure 6.**
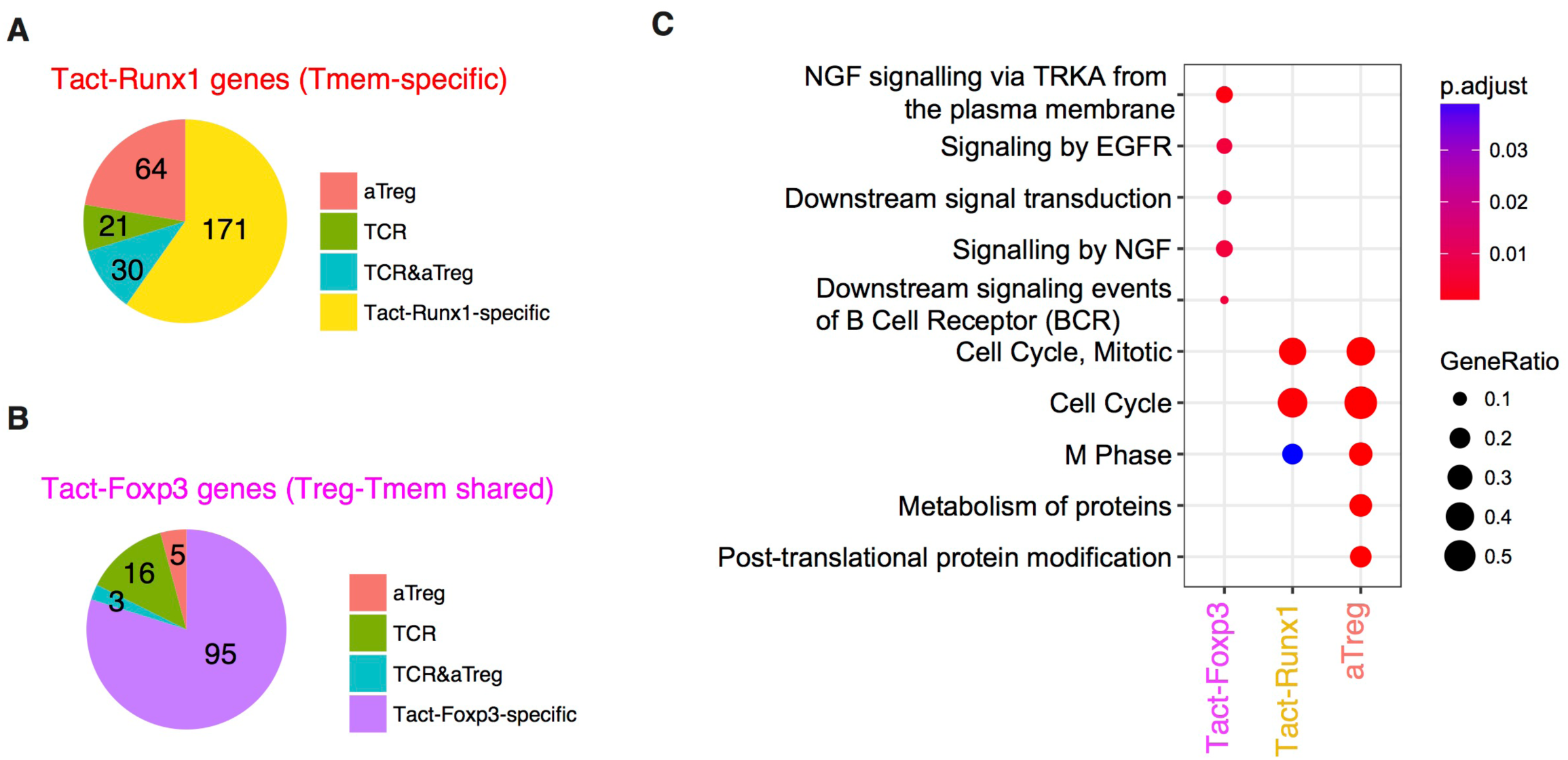
The comparative analysis of Tmem-specific and Treg-Tmem shared activation genes and TCR-dependent and activated Treg-specific genes. Venn diagram analysis was used to obtain intersects of *TCR-dependent genes* (DEG between TCRa KO and WT Treg), *aTreg-specific genes* (DEG between aTreg and rTreg), and Tact-Foxp3 and Tact-Runx1 genes (see Figure 3). (**A**) Pie chart showing the number of genes in the intersects between *aTreg-specific genes*, *TCR-dependent genes,* and Tact-Runx1 genes. (**B**) Pie chart showing the numbers of *aTreg-specific genes*, *TCR- dependent genes,* and Tact-Foxp3 genes. (**C**) Pathway analysis of Tact-Foxp3 genes, Tact-Runx1 genes and *aTreg-specific genes* showing enriched pathways in these gene lists.

### FOXP3 expression more frequently occurs in activated T cells than resting cells by single cell CCA

The analyses above showed that Treg are on average more activated than naïve T cells and that the activation status of Treg can be variable. However, it is still unclear whether individual Treg are activated than any naïve T cells at the single cell level. The alternative hypothesis is that Treg are enriched with the T cells that have recognised their cognate antigens and been activated. In order to determine this and thereby understand the dynamics of T cell regulation in vivo, we investigated the single cell RNA-seq data of tumour-infiltrating T cells from human patients (Ref. (35) **Table 1**), and further enquired how the activation mechanisms are operating in Treg at the single cell level.

Firstly, we *in silico*-sorted FOXP3+ and FOXP3- CD4^+^CD3^+^ T cells from unannotated single cell data from tumours, which tissues were dispersed and CD45+ cells were sorted by flow cytometry without the use of any lymphocyte markers (GSE72056, **Table 1**). Thus, the identities of individual single cells were needed to be identified in a data-oriented manner, and Treg and non-Treg cells in these tumour tissues had unknown individual activation and differentiation statuses. Thus, we applied CCA to the *in silico*-sorted single cell T cell data using the explanatory variables of activated conventional CD4+ T cells (*Tact*) and resting T cells (*Trest*; GSE15390, **Table 1**), aiming to define individual single cells according to their level of activation by their correlations to these two variables (**Figure 7A**). Here we used these two variables, *Tact* and *Trest*, in order to generate a two-dimensional CCA solution, instead of a single explanatory variable that represents T cell activation by the log2 fold change in gene expression between activated and resting CD4+ T cells (*c.f.* Figures 1-6), which produces a one-dimensional CCA solution visualised as a single axis), because we aimed to identify any additional major differentiation process(es) in the Axis 2. The explanatory variables *Tact* and *Trest* are both captured by Axis 1 because they represent two poles of one continuum – the spectrum of activation – ranging from ‘resting’ to ‘activated’ cell state. Thus the CCA aimed to sort the single cells according to their individual levels of activation along the spectrum of activation, capturing the heterogeneity in activation levels in single cells. Remarkably, in the cell sample space of the CCA solution, the majority of FOXP3+ T cells were positively correlated with the T cell activation variable *Tact* (**Figure 7B**), and thus had negative scores in the Axis 1 (**Figure 7B**). Here, CCA Axis 1 *×* (−1) score is designated as the T cell activation score. Thus, using the activation score and FOXP3 expression, the following four subpopulations were defined: “Activated FOXP3+”, “Resting FOXP3+”, “Activated FOXP3-”, and “Resting FOXP3-” (**Figure 7B**).

**Figure 7.**
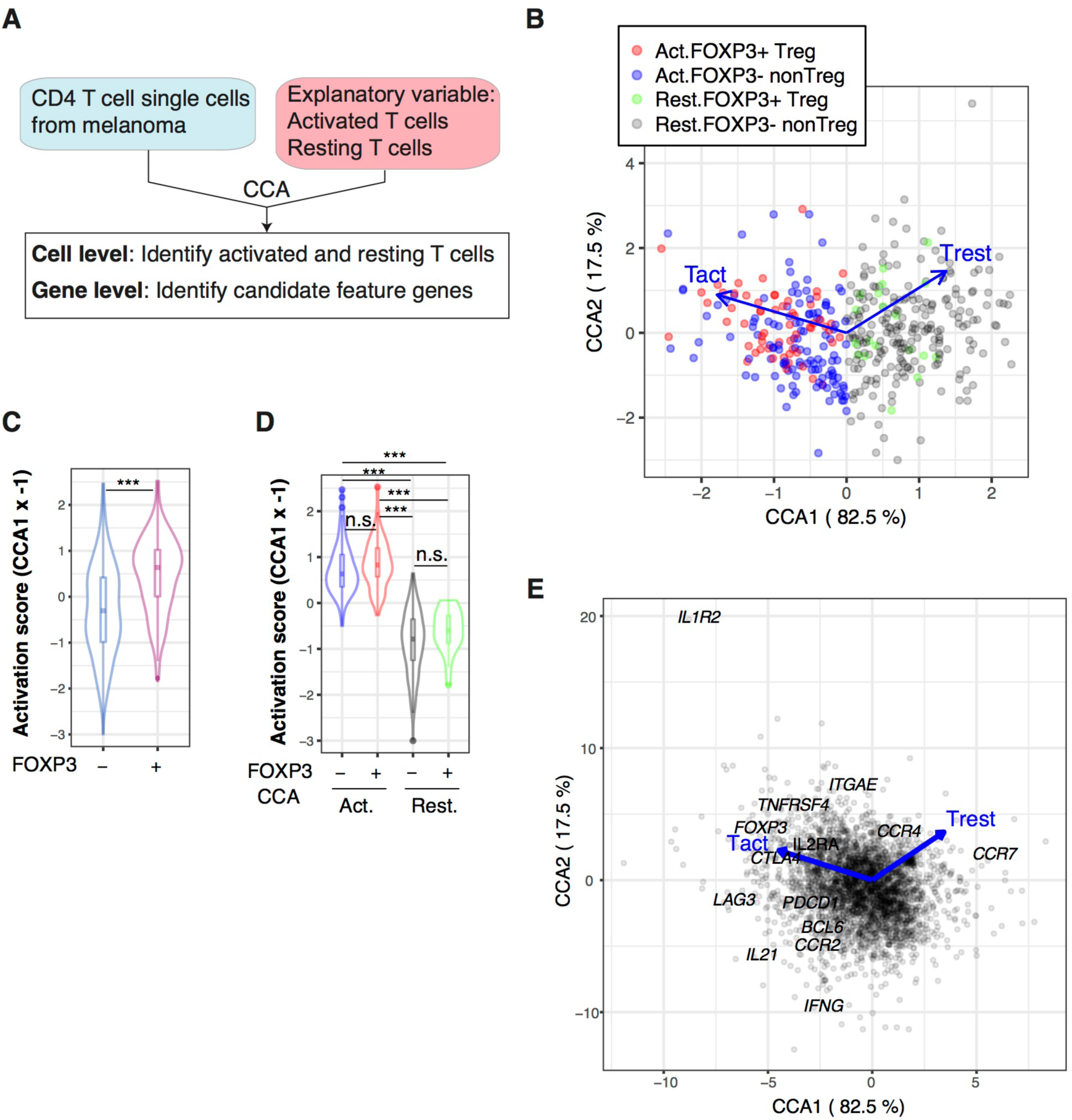
Single cell CCA of melanoma-infiltrating T cells determines the activation status of individual T cells and identifies a putative Tfh-like process. (**A**) Schematic representation of CCA of CD4+ T cell single cell transcriptomes analysed by two explanatory variables: activated naïve T cells (Tact) and resting naïve T cells (Trest). (**B**) CCA biplot showing the relationship between Treg and non-Treg T cells (sample scores) and the explanatory variables (Tact and Trest). Axis 1 represents the difference between Tact and Trest, and thus, Activated T cells and Resting T cells were defined by the CCA Axis 1 score, and these cells were further classified into Treg and non-Treg by their *FOXP3* expression (see legend). Percentage indicates that of the variance (inertia) accounted for by the axis. (**C**) Violin plot showing the CCA activation scores (Axis 1 score *×* −1) of FOXP3- and FOXP3+ cell groups. Asterisk indicates statistical significance by Mann-Whitney test (**D**) Violin plot showing the CCA activation scores of Activated (Act.) and Resting (Rest.) FOXP3- and FOXP3+ cell groups. Asterisks indicate the values of post-hoc Dunn’s test following a Kruskal Wallis test. *** p < 0.005. (**E**) Gene biplot of the CCA solution in (B) showing the relationships between genes (grey circles) and the Tact and Trest explanatory variables (blue arrows). Genes are shown by grey circles, and well-known genes that are key for T cell activation processes are annotated.

Next, we aimed to determine whether individual activated FOXP3+ Treg are more activated than activated FOXP3- non-Treg at the single cell level According to the T cell activation score established by the CCA solution in Figure 7B, FOXP3^+^ Treg had significantly higher T cell activation scores than FOXP3- non-Treg on average, as indicated by the higher median in the violin plots and greater density of samples with higher T cell activation scores (**Figure 7C**), confirming the results by bulk cell analysis (Figure 1). Using the CCA definition of activated and resting Treg and non-Treg established in Figure 7B, the T cell activation score neatly captured the activated status of single cells, allocating high positive and negative scores to activated and resting cells, respectively (**Figure 7D**). Importantly, there was no significant difference between Activated FOXP3+ and Activated FOXP3- cells and between Resting FOXP3+ and Resting FOXP3- cells (**Figure 7D**), indicating that in tumour microenvironment, Treg cells are as activated as non-Treg CD4+ T cells, which may be enriched with Teff. Strikingly, 32.5% of activated T cells expressed FOXP3, while only 8.2% of resting T cells expressed FOXP3 in Figure 7B. In other words, FOXP3 expression occurred more frequently in activated T cells. Given that the activation signature of Treg is dependent on TCR signals (Figure 5), these results suggest that FOXP3 expression occurs predominantly in the activated T cells that have recognised the tumour antigens and received TCR signals, as a negative feedback mechanism to suppress the effector response against tumour antigen (7). Alternatively, but not exclusively, FOXP3+ T cells may have high-affinity TCRs to self-MHC and/or tumour antigens and be more prone to activation (10).

In the gene space of the CCA solution, genes with strong correlations to activated FOXP3^+^ T cells included *FOXP3* itself and common Treg markers such as *CTLA4* and *IL2RA* (CD25), which were found in the upper left quadrant (Axis 1-negative Axis 2-positive). Interestingly, the lower left quadrant (Axis 1-negative Axis 2-negative) contained more Tfh-like or effector-like molecules *PDCD1* (PD-1), *BCL6*, *IL21*, and *IFNG.* The chemokine receptors *CCR5* and *CCR2* had negative scores in Axis 1 (i.e. correlated with *Tact*), while *CCR7* had a high positive score in Axis 1 (i.e. correlated with *Trest*) (**Figure 7E**).

### Identification of Tfh-like differentiation and Foxp3-driven processes and the common activation process in tumour-infiltrating T cells

Next, we aimed to identify major differentiation and activation processes in the single cell transcriptomes above. To this end, we have developed a new CCA approach for single cell analysis (Single Cell Combinatorial CCA, SC4A), which aims to visualise major differentiation/activation processes and the underlying gene regulations (**Figure 8A**, see Materials and Methods). Firstly, we classified single cells into the four populations (Activated and Resting cells, and FOXP3+ Treg and FOXP3- non-Treg; Figure 7B), and thereby identified the following four processes as putative differentiation and activation processes in the dataset: T cell activation (Activated cells), and naïve-ness (Resting cells), FOXP3-driven process (Activated FOXP3+), and Tfh-like process (Activated FOXP3-) (Figure 7). Secondly, based on their high scores in the CCA solution (i.e. either high positive or high negative scores in either Axis 1 or 2 in Figure 7E) and abundant expressions in FOXP3+ and FOXP3- cells (data not shown), we selected 12 candidate genes (*CCR7*, *CCR5*, *CCR4*, *IL2RA*, *IL2RB*, *CTLA4*, *ICOS*, *TNFRSF4*, *TNFRSF9*, *FOXP3*, *BCL6*, *PDCD1*) as the candidate genes for the four processes. From these genes, we identified the most positively correlated gene to each of the four processes using the combinatorial CCA, which tests all the combinations of variables by CCA and obtains the most correlated gene for each population; see Materials and Methods). Thus, *PDCD1, FOXP3*, *CTLA4*, and *CCR7* were identified as the most correlated genes for Activated FOXP3-, Activated FOXP3+, Activated T cells, and Resting T cells, respectively (**Supplementary Figure 3**), which represent the four immunological processes (see above). Finally, using these four genes as explanatory variables, we applied CCA to the single cell transcriptomes, obtaining the solution of the SC4A approach.

The single cell space of the SC4A solution showed that Activated and Resting T cells had negative and positive scores, respectively (**Figure 8B**). This indicates that Axis 1 represents T cell activation vs naïve-ness. Single cells were successfully clustered into Activated FOXP3^+^ Treg, Activated FOXP3- non-Treg, and Resting T cells. Resting FOXP3+ Treg and Resting FOXP3- T cells were mostly overlapped (**Figure 8C**), indicating that the major features in the dataset dominated the difference between these two resting T cell groups. Importantly, the explanatory variable CTLA4, which represents the T cell activation process, was highly correlated with both Activated FOXP3^+^ Treg and Activated FOXP3- non-Treg at the middle, indicating its neutral position in terms of Tfh and Treg activation processes. As expected, the variable CCR7, which represents naïve-ness, was correlated with both Resting FOXP3+ Treg and Resting FOXP3- T cells. The explanatory variable PDCD1, which represents the Tfh-like process, was highly correlated with Activated FOXP3- non-Treg cells, while the variable FOXP3 was correlated with Activated FOXP3+ Treg. Thus, the single cell transcriptomes were modelled by the correlations between gene expression, single cells, and the expression of the four key genes, which represent the four immunological processes (**Figure 8B and 8C**). PCA and t-distributed stochastic neighbor embedding (t-SNE) did not provide insights into such cross-level relationships or clear separations of the populations (**Supplementary Figure 4**).

Next, in order to understand the relationship between the T cell activation signature and FOXP3-driven and Tfh-like processes (Figure 8B and 8C), we aimed to identify and characterise genes with high correlations to these processes, which were represented by CTLA4, FOXP3, and PDCD1 explanatory variables, by analysing the gene space of the final output of SC4A (Figure 8C; see Methods). As expected, the Tfh genes, *IL21* and *BCL6* (45), were highly correlated with PDCD1 explanatory variable. *IL2RA* (CD25) is a Treg marker (46) and was highly correlated with FOXP3 explanatory variable. *IL7R* and *BACH2* are known to be associated with naïve T cells (47, 48), and were positively correlated with CCR7 explanatory variable, which represents the naïve-ness (**Figure 8C**). Thus we defined *FOXP3-driven Treg genes* (magenta circles) and *Tfh-like genes* (blue circles) according to their high correlation to the FOXP3 and the PDCD1 explanatory variables, respectively, while we designated as *Activation genes* (red circles) the genes that have high correlations with the CTLA4 variable, including *LAG3* and *CCR5*, which were positioned around 0 in Axis 2 (Figure 8C).

### Identification of the bifurcation point of activated T cells that leads to Tfh-like and Treg differentiation in tumour-infiltrating T cells

The analyses above strongly suggested that there are two major differentiation pathways for those tumour-infiltrating T cells, which are regulated by FOXP3-driven and Tfh-like processes. In order to identify these lineages, we applied an unsupervised clustering algorithm to the sample space of the SC4A/CCA result (Figure 8B), and identified 6 clusters, to which a pseudotime method (49) was applied, constructing “lineage curves” (**Figure 8D**; see **Methods**). Importantly, the lineage curves had a bifurcation point at Cluster II, which leads to the two distinct differentiation pathways, Tfh-like and FOXP3-driven differentiation. Since cells may change and mature their phenotypes in different dynamics between these two lineages, we designated Tfh-like-associated and FOXP3-associated pseudotime as Tfh-pseudotime and FOXP3-pseudotime (**Figure 8D**).

In fact, the expression of *Activation genes* was progressively increased in the shared clusters (i.e. Cluster I and II) for the two pseudotimes, and throughout the rest of the FOXP3-pseudotime and the early phase of Tfh-like differentiation (i.e. Cluster III) in Tfh-pseudotime, while it was suppressed towards the end of Tfh-like differentiation (Cluster IV; **Figure 8E**) in Tfh-pseudotime. Given that Tfh-pseudotime is correlated with *PDCD1* expression (**Figure 8C**), this suggests that *PDCD1* expression and the Tfh-effector process are induced during the earlier phases of effector T cell activity, and that the activation processes in *PDCD1*^high^ T cells are suppressed, presumably through PD1-PDL1 interactions in the tumour environment (50). Interestingly, *FOXP3-driven genes* had similar dynamics to *Activation genes* in both FOXP3-pseudotime and Tfh-pseudotime (**Figure 8F**). In contrast, *Tfh-like genes* were mostly suppressed throughout FOXP3-pseudotime, while they were progressively induced throughout Tfh-pseudotime (**Figure 8G**). These differential regulations of two gene modules resonate with those of Tact-Foxp3 genes (which are expressed by both Treg and Tmem) and Tact-Runx1 genes (which are expressed specifically in Tmem, and repressed in Treg) (Figure 3). In fact, *FOXP3* expression is weakly induced in some cells in the bifurcating Cluster II and the early phase of Tfh-like differentiation (Cluster III) in Tfh-pseudotime, and is progressively increased at and beyond Cluster V in FOXP3-pseudotime (**Figure 8H**).

*RUNX1* is highly expressed in the common Clusters I and II, and is downregulated in the transition from Cluster II to Cluster III in Tfh-pseudotime, and from Cluster II to Cluster V in FOXP3-pseudotime (**Figure 8I**), which is compatible with the known dynamics of RUNX1 expression: Runx1 is downregulated when naïve CD4+ T cells differentiate into activated/effector cells following TCR signaling (51). By analysing other key genes used as CCA explanatory variables, *CTLA4* was induced at the bifurcating point, Cluster II, and onwards in both of the lineages at equivalent expression levels (**Figure 8J**), reflecting the activated status of both effector Tfh-type cells and Treg. Importantly, CTLA4 is a marker of Treg as well as activated effector T cells, where it acts as a negative regulator of T cell proliferation (52).

*PDCD1* expression was also induced at the bifurcating point, and throughout Tfh-pseudotime, but specifically suppressed in the early phase of FOXP3-pseudotime (**Figure 8K**), which is compatible with the known dynamics that PDCD1 is transiently upregulated in activated CD4+ T cells as a negative regulatory mechanism to restrain proinflammatory immune responses and maintain peripheral tolerance (53). Further supporting this dynamic perspective, *IL2* expression occurs mainly in Cluster II, indicating that these cells are enriched with the T cells that recently recognised antigens (54) (**Figure 8L**). Consistently, the expression of the naïve T cell marker CCR7 was the highest in the cells with a relatively naïve phenotype in shared Cluster I, and was moderately downregulated in the early and late phase of Tfh-pseudotime, and suppressed in most Treg in FOXP3-pseudotime (**Figure 8M**).

These results collectively support the model that constant activation processes in the tumour microenvironment promote terminal differentiation of the Treg- and Tfh-like lineages in both previously committed and non-committed lineages of T cells. Interestingly, Cluster II is the bifurcation point, in which T cells show moderate activation and together with simultaneous expression of FOXP3 and Tfh-like genes, as well as RUNX1 and PDCD1 expression. These cells are most probably engaged in decision-making about their cell fate and the cell type-specific usage of these genes – whether their transcriptional mechanisms would be used to generate a proinflamamtory or regulatory response. This understanding was possible because SC4A effectively annotated genes and cells and thereby allowed to identify new cell populations.

### Identification of markers for the differential regulation of Tfh-like and Treg differentiation in activated T cells

Lastly, we aimed to demonstrate the utility of the current approach by discovering exemplary marker genes that distinguish cells in FOXP3- and Tfh-pseudotime (i.e. the FOXP3-driven pathway I-II-V-VI, and the Tfh-like pathway I-II-III-IV) (**Figure 9A**), and identifying the T cell subpopulations by a flow cytometric visualisation of single cell data. Since *Activation genes* (Figure 8C) are shared by early phases of Tfh-like and FOXP3-driven differentiation (Figure 8E), we took the intersect of these genes and the Tact-Foxp3 genes, which were expressed by both resting Tmem and resting Treg in mice (Figure 3). *DUSP4* and *NFAT5* were such genes and in fact induced in cells at the activated bifurcating Cluster II and onwards in both lineages (**Figure 9B**). Similarly, in order to identify a marker to distinguish Treg and Tfh-like cells, firstly, we identified *CCR8* and *IL2RA* in the intersect of FOXP3-driven genes (Figure 8C) and the Tact-Foxp3 genes, which were induced highly and progressively in Treg-lineage cells throughout FOXP3-pseudotime, while mostly suppressed across Tfh-pseudotime (**Figure 9C**). In contrast, *BCL6* and *KCNK5* (found in the intersect of Tfh-like genes (Figure 8C) and the Tact-Runx1 genes, which are expressed in resting Tmem but suppressed in resting Treg (Figure 3)) were progressively induced across Tfh-pseudotime, while suppressed in FOXP3-pseudotime (**Figure 9D**).

**Figure 8.**
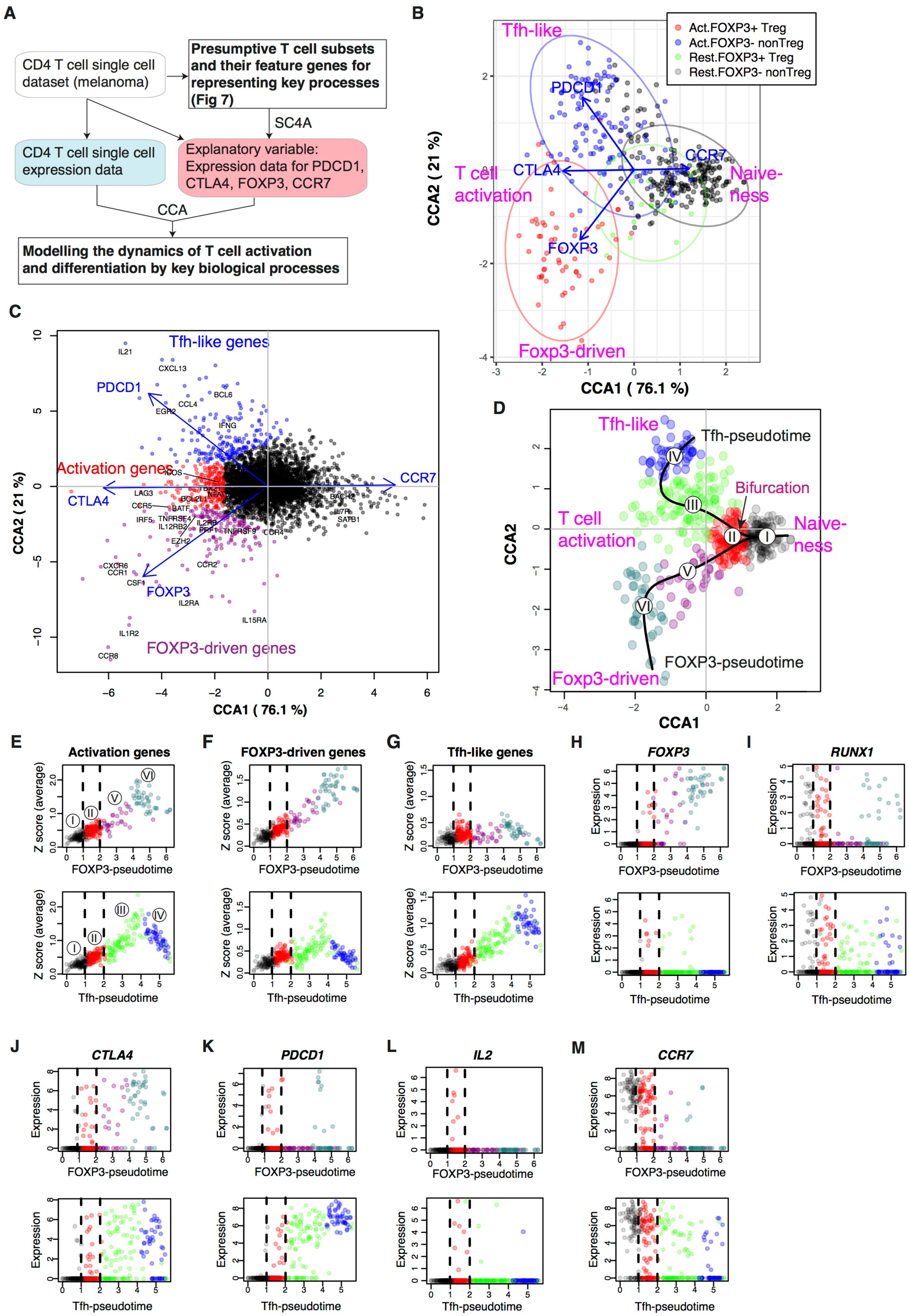
SC4A identifies the bifurcation point of activated T cells that leads to Tfh-like and Treg differentiation in tumour-infiltrating T cells. SC4A was applied to the single cell data of tumour-infiltrating T cells, and 4 genes (CTLA-4, CCR7, FOXP3, and PDCD1) were chosen as explanatory variables to represent the T cell activation, resting, FOXP3-driven process, and Tfh-like process. **(A)** The design of analysis. The single cell data from the melanoma samples were analysed by SC4A to identify the most effective combinations of explanatory variables for dispersing the 4 presumptive T cell populations identified in Figure 7. These genes were used as explanatory variables to analyse the rest of the single cell data as main dataset. Thus, the single-cell level dynamics of T cell differentiation and activation are modelled by the key biological processes that are represented by the T cell populations and explanatory variables. (**B**) Single cell sample space of the final SC4A output showing correlations between single cell samples and the explanatory variables **(C)** Gene space of the final SC4A output showing correlations between genes and the explanatory variables. The genes that showed high correlations to the PDCD1, CTLA4, and FOXP3 variables were identified as Tfh-like genes, Activation genes, and FOXP3-driven genes, respectively. **(D)** The identification of two differentiation processes as lineages and a bifurcation point. The cells in the sample space of the SC4A output (B) were classified into 6 clusters by an unsupervised clustering algorithm. These clusters were further analysed for pseudotime inference. (**E-G**) The average gene expression was plotted against each pseudotime (upper: FOXP3- pseudotime; lower: Tfh-pseudotime). The bifurcation point (Cluster II) is emphasised by broken lines. The numbers in circle indicate the cluster number. Gene expression was standardised, and the sum of the standardised expression was obtained for (**E**) Activation genes, (**F**) FOXP3-driven genes, and (**G**) Tfh-like genes (see C). **(H-M)** The expression of key genes was plotted against each pseudotime.

**Figure 9.**
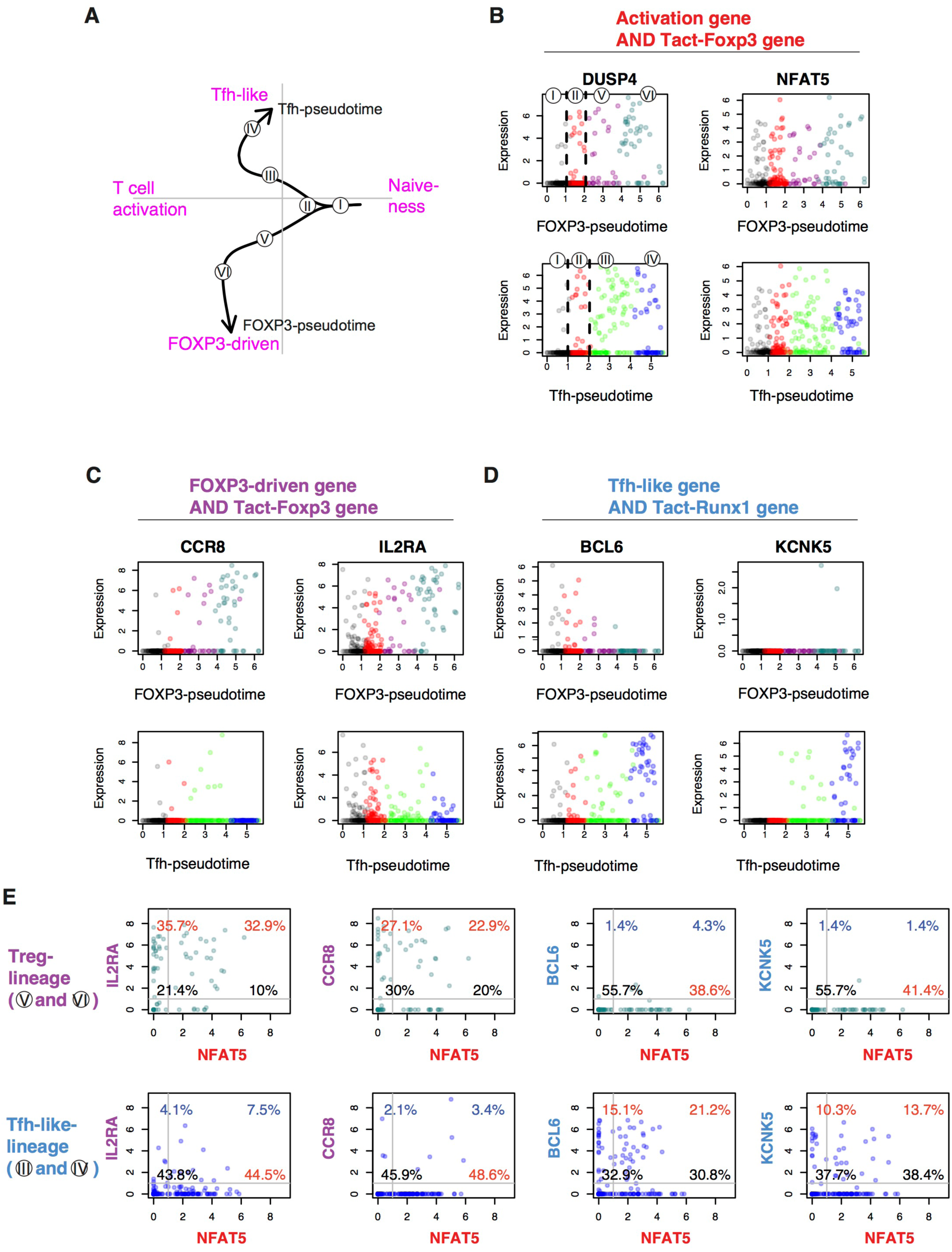
Identification of the conserved genes for the differential regulation of Tfh-like and Treg differentiation in activated T cells. **(A)** The identified lineage curves and the bifurcation point in the tumour-infiltrating T cells. The number in circle indicates the cluster number in Figure 8D. **(B-D)** The expression of selected feature genes was plotted against each pseudotime. Genes are from the intersect of **(B)** Activation genes (Figure 8C) and Tact-Foxp3 genes (Figure 3), **(C)** FOXP3-driven genes (Figure 8C) and Tact-Foxp3 genes, and **(D)** Tfh-like genes (Figure 8C) and Tact-Runx1 genes (Figure 3). **(E)** The expression of selected genes in the tumour-infiltrating T cells was shown by a 2-dimensional plot in a flow cytometric style. Data from Treg-lineage cells (Cluster V and VI, upper panels) and Tfh-like lineage cells (Cluster III and IV, lower panels). The gene in x-axis (NFAT5) is from the activation gene group (B), while y-axis shows genes from either the FOXP3- Treg group (C) or the Tfh-like/Tmem group (D). Thresholds and quadrant gates were determined in an empirical manner using density plot.

Lastly, in order to make the newly obtained knowledge easily accessible to experimental immunologists, we showed the expression of *NFAT5*, *IL2RA*, *CCR8*, *BCL6*, and *KCNK5* in the tumour-infiltrating T cells in a flow cytometric format (**Figure 9E**). The common activation gene *NFAT5* in fact captured the majority of Treg-lineage cells (i.e. cells in the Clusters V and VI) and Tfh-like-lineage cells (i.e. cells in the Clusters III and IV). The Treg-specific genes the expression of *IL2RA* and *CCR8* occurred in the majority of FOXP3+ Treg-lineage cells, whether NFAT-positive or negative, but not in most of Tfh-like-lineage cells. In contrast, the Tfh-like-specific genes *BCL6* and *KCNK5* were expressed by a majority of Tfh-like-lineage cells and were not expressed in Treg-lineage cells (**Figure 9E**).

Collectively, these results indicate that the SC4A analysis successfully decomposed the gene regulations for T cell activation and Treg and effector T cell differentiation, identifying new cell populations, which include activated cells at the bifurcation point, early and late phases of Treg and Tfh-like differentiation, and their feature genes. In addition, although there must be considerable differences between resting T cells in the secondary lymphoid organs and between humans and mice, our study successfully identified the shared activation processes and the conserved genes that are differentially used between the Treg- and the Teff-lineage cells, identifying a shared systems-level mechanism for the differentiation regulation of activation and differentiation processes in CD4+ T cell populations.

## Discussion

Resting Treg showed an activated status, comparable to that of Teff and Tmem at the population level. In addition, the activation signature of Treg was more remarkable in CD44^hi^CD62L^lo^ activated Treg than CD44^lo^CD62L^hi^ naïve-like Treg. CD44^hi^CD62L^lo^ Treg are also identified as eTreg, which may have enhanced immunosuppressive activities (55). The eTreg fraction includes the GITR^hi^PD-1^hi^CD25^hi^ “Triple-high” eTreg that have high CD5 and Nur77 expressions, which indicates that they have received strong TCR signals (17). In humans, CD25^hi^CD45RA*^−^*FOXP3^hi^ eTreg highly express Ki67 (56), indicating that these cells were recently activated. Given that TCRs of Treg have higher affinities to self-antigens (57), these eTreg may have the most self-reactive TCRs during homeostasis. Alternatively, the eTreg subset may have recently received strong TCR signals and upregulated activation markers, and such cells may acquire a resting status at later time points. Future investigations by TCR repertoire analysis will answer this question.

Our study revealed the heterogeneity of FOXP3+ Treg at the single cell level, and showed that tumour-infiltrating Treg include FOXP3+ T cells with various levels of activation (Figure 7B and Figure 8C). It is plausible that, in the physiological polyclonal settings, the variations in the activated status of individual Treg may be due to the TCR affinity to its cognate antigen, the availability of cognate antigen, and the strength and duration of TCR signals. Our SC4A analysis identified the *FOXP3*-driven genes, which are specific to activated FOXP3+ cells and include IL-2 and common gamma chain cytokine receptors (i.e. *IL2RA*, *IL2RB*, *IL15RA*, *IL4R*, and *IL2RG*), DNA replication licensing factors (e.g. MCM2), and transcription factors such as *PRDM1* (BLIMP1) and *IRF4* (which control the differentiation and function of eTreg (19)). These gene modules are distinct from the Tfh-like genes and the activation genes (Figure 8), and may be controlled specifically by *FOXP3* under strong TCR signals. The expression of these genes is variable within the FOXP3+ T cells, suggesting that the transcriptional activities of these genes are dynamically regulated over time in tumour-infiltrating Treg. Thus, single cell-level analysis is becoming a key technology to address the heterogeneity of Treg. To our knowledge, this study is one of the first single cell analyses of Treg transcriptomes, while we find that, during the review process of this manuscript, another study addressing Treg heterogeneity by single cell RNA-seq was deposited at a preprint server (58)).

The shared activation genes between activated FOXP3+ Treg and FOXP3- non-Treg contain apoptosis-related genes (e.g. *CASP3*, *BAD*), which may be differentially controlled between Treg and non-Treg at the protein level. For example, activated FOXP3- non-Treg express *DUSP6* (Figure 9B), which is a negative regulator of JNK-induced apoptosis through BIM activation, while *FOXP3* suppresses *DUSP6* expression and promotes the apoptosis mechanism (59). In addition, the activation genes include transcription factors such as *TBX21* (T-bet) and *BATF*. Although *TBX21* is sometimes thought to be a Th1-specific gene, it is upregulated immediately after T cell activation (60). *BATF* was identified as a critical factor for the differentiation and accumulation of tissue-infiltrating Treg (61). These activation genes may be required when T cells are activated and differentiate into either Treg or Teff. Further studies are required to investigate the temporal sequences of these differentiation events *in vivo*.

Although the effects of TCR signals on Tmem were not directly examined, considering that Tmem are self-reactive and their differentiation is dependent on the recognition of cognate antigens in the thymus (7), these results collectively suggest that the activation signature of Tmem is also dependent on TCR signals, as is the activation signature in Treg (Figure 5B). Intriguingly, some Treg may lose their Foxp3 expression and become ex-Treg, which are enriched in CD44^hi^ effector T cells or Tmem (30). In contrast, a Tmem population (precisely, Foxp3^−^CD44^hi^CD73^hi^FR4^hi^ T cells) efficiently express Foxp3 during lymphopenia (62). These findings support the feedback control model that Foxp3 expression can be induced in Tmem and sustained in Treg as a regulatory feedback mechanism for TCR signals (7). Given the variations in the activated status in individual Treg and Tmem, single cell analysis will be required to address this problem. For example, although Samstein *et al.* showed that DNA hypersensitivity sites in Treg are similar to those in activated T cells (9), it is possible that DNA hypersensitivity sites are variable between individual Treg, and that Tmem may have a similar chromatin structure to Treg.

Importantly, our analysis showed that Tmem-specific activation-induced genes (i.e. Tact-Runx1 genes) are uniquely repressed in Treg. The repression is likely to be mediated by the interaction between Foxp3 and other transcription factors that regulate the expression of the Tmem-specific activation genes (Figure 3C). Interestingly, *Runx1* was associated with these Tmem-specific genes. In fact, Foxp3 interacts with Runx1 and thereby represses IL-2 transcription and controls the regulatory function of Treg (38), and a significant part of the Foxp3-binding to active enhancers occurs through the Foxp3-Runx1 interaction (9). These suggest that Runx1 may have a unique role in the differentiation and maintenance of Tmem.

While CTLA-4 is commonly recognised as a Treg marker, it is upregulated in all activated T cells, thus CTLA-4 is also a marker of activated T cells (41). CTLA-4 is in fact expressed by only a subset of resting Treg (63), which may be more activated and proliferating *in vivo* (64). In fact, our study shows that CTLA-4 is expressed by non-Treg activated T cells including resting Tmem (Figure 3D) and FOXP3- Tfh-like effector T cells in the tumour microenvironment (Figure 7E and 8C). These findings support that CTLA-4 is primarily a marker for general T cell activation, rather than Treg-specific marker, and that Treg are highly activated T cells with *FOXP3* and *CTLA-4* expression. Importantly, although both FOXP3+ and FOXP3- cells had the same relative level of activation (Figure 7D), the absolute number of FOXP3+ cells expressing *CTLA4* was lower than that of Tfh-type cells (Figure 8J), which suggests that therapeutic anti-CTLA4 antibodies (i.e. Ipilimumab) primarily target activated Tfh-like effector cells and thereby directly enhance their activities in tumour microenvironments. Future studies are required to experimentally investigate the *in vivo* dynamics of CTLA-4 expression in mice and humans.

In contrast, the expression of *PDCD1* was consistently high in all Tfh-like cells, while it was sparse among FOXP3+ cells (Figure 8K). The co-expression of *BCL6* and *IL21* in some of these PD-1+ cells indicates that Tfh differentiation occurs in the tumour microenvironment, presumably through the repeated and chronic exposure to quasi-self antigens (i.e. tumour antigens). Interestingly, the Tfh signature has been identified in type-I diabetes in both mice and humans (65). Intriguingly, the Tfh-like genes include cell-cycle related genes (e.g. *CDK6*), immediate early transcription factors (*NFATC1*, *EGR2/3*), and RNA-processing genes (e.g. *DICER1*). The significance of these gene modules should be addressed in future studies. However, the high *PDCD1* expression in Tfh-like cells may make them vulnerable to the negative immunoregulatory effect of PD-1 in tumour microenvironments (50). In fact, the most mature *PDCD1*^high^ Tfh-like cells (cluster VI, Figure 8K) moderately decrease the expression levels of activation genes (Figure 8E), suggesting that these cells may have started to be regulated by PD1 ligands. Further experimental investigations are required to reveal how dynamically PD1 regulates T cells during immune response.

Interestingly, *RUNX1* is completely repressed in the early phase of FOXP3- pseudotime (Cluster V) but re-expressed in the late phase of FOXP3- pseudotime the expression of *RUNX1* is significantly elevated (Cluster VI) (**Figure 8I**). Similar to RUNX1, some cells appear to be expressing *PDCD1* in the later phase of FOXP3-pseudotime in Treg-lineage cells. The reappearance of effector phenotype genes *RUNX1* and *PDCD1* in FOXP3^high^ cells may indicate that these Treg are highly activated effector Treg, which may be actively participating in neutralizing the activities of effector T cells and Tfh. Alternatively, these FOXP3^high^ cells may include ambivalent cells with the characters of both regulatory and effector, or alternatively, be at the point of conversion to Tmem (**Figure 8K**). In bulk resting Treg, *Pdcd1* was expressed at low levels in some Treg (Figure 3E) as well as PD-L2-encoding gene *Pdcd1lg2* (Figure 3D). Future studies are required to reveal the role of these cells.

SC4A is a useful method to identify distinct clusters of T cells and the correlated genes to each cluster, and thereby to reveal characteristic cell groups and their active gene modules, while retaining the single-cell level variations. We also showed that SC4A and CCA results can be further analysed by the *pseudotime* approach (CCA-pseudotime). Since SC4A/CCA provides functional annotations to cell groups and gene clusters, the understanding of the pseudotime axis is effective, as shown in the current study. However, given that pseudotime is not a direct measurement of the time-dependent events, but rather is that of similarities between samples (66), future studies are required to analyse time-dependent events *in vivo,* ideally with a new experimental system to directly address the temporal dynamics. In order to make the current SC4A/CCA approach accessible to experimental immunologists, we visualised our single cell data findings using a flow-cytometry style (Figure 9). Although reliable antibodies are currently not available for those intracellular candidate genes, and the expression of protein and transcripts may not be synchronized, the present study demonstrated the power of the SC4A/CCA approach to extract biological meaning from unannotated single cell RNA-seq data. The current limitation of SC4A is that it is computationally expensive (i.e. requires several hours for each analysis using a standard desktop), and the improvement of the computational algorithm using a low-level language will be beneficial. Importantly, SC4A is most effective when used together with in-depth knowledge of immunology and gene regulation, facilitates the interpretation of CCA results and explanatory variable selection. Thus, it is hoped that these tools will be widely used by experimental immunologists with a sound understanding of the biological significance of results, as well as adequate competence in computational analysis, which will enable to ask questions involving multidimensional problems such as multiple T cell subsets.

## Data and code availability

All R codes are available upon request. Processed data will be provided upon reasonable requests to the corresponding author.

## Acknowledgement

We thank Dr Cristina Leslie for providing us the details of their dataset GSE83315, Prof Lucy Walker for a useful discussion, and Dr David Bending for valuable comments on the manuscript. M.O. is a David Phillips Fellow (BB/J013951/2) from the Biotechnology and Biological Sciences Research Council (BBSRC), and is also supported by a pump-priming grant from Cancer Research Centre of Excellence, Imperial College London/the Institute of Cancer Research. A.B is supported by the Japanese Ministry of Education, Culture, Sports, Science and Technology (MEXT) PhD scholarship.

